# Host-derived bile acids drive dysbiosis by selecting bile-resistant epimerizing bacteria in inflammatory bowel disease

**DOI:** 10.64898/2026.06.18.733240

**Authors:** Maximilian Baumgartner, Franziska Schnaufer, Maeva Duquesnoy, Takahiro Asatsuma, Adrija Chakrabarty, Adrian Frick, Claudia Fuchs, Michael Gerstorfer, Patrik Hains, Thomas Köcher, Patrick Schimmel, Mauriz A. Lichtenstein, Sofia Leistl, Felix Pinter, Elena Krstevska, Thet Khaing Nyein, Christoph Högenauer, Athanasios Makristathis, Christoph Gasche, Christian Primas, Walter Reinisch, Georg E. Winter, Michael Trauner, Claudia Günther, Georg Busslinger, Gregor Gorkiewicz, Benoit Chassaing, Clarissa Campbell

**Author notes:** Contributed equally (Franziska Schnaufer, Maeva Duquesnoy).

## Abstract

Microbial dysbiosis is a hallmark of inflammatory bowel diseases (IBD); however, its drivers and impact on disease pathophysiology are poorly understood. Applying neural network-based feature attribution to metabolomics and metagenomics datasets from >5000 individuals, we identified epimerized host derived bile acids (BAs) produced by microbial hydroxysteroid dehydrogenases (HSDHs) as a novel hallmark of IBD-associated dysbiosis. Epimerized BAs reduce FXR activity in intestinal epithelial cells and dampen their production of FGF19, a negative feedback regulator of host-derived bile acid (HBA) production in the liver. Increased HBA levels drive colonic epithelial remodeling by impacting goblet cell maturation and select for HSDH-carrying bacteria that transform bactericidal HBA into less toxic, epimerized forms. Confirming the translational relevance of these findings, we demonstrated that high HBA levels limit fecal microbiota transplant engraftment and show that BA sequestering drugs support microbiome recovery in patients with high HBA levels. Together, we discover that elevated HBAs deplete BA-sensitive commensals and favor the growth of HSDH-encoding pathobionts that disrupt host BA feedback signaling, establishing a causal link between changes in microbial ecology and IBD pathophysiology.

## Introduction

Inflammatory bowel diseases (IBD) including Crohn’s disease (CD) and ulcerative colitis (UC) are chronic, increasingly prevalent disorders with complex, multifactorial origins^1^. Despite advances in immunomodulatory therapies, many patients do not achieve sustained remission, reflecting an incomplete understanding of disease mechanisms^2,3^. Growing evidence supports the interplay between the intestinal microbiome, host metabolism, and immune responses as central to disease onset and progression^4,5^. While dysbiosis is consistently associated with IBD^6^, its ecological drivers and the mechanisms by which the microbiome influences IBD pathophysiology remain poorly understood^7^.

Perturbations in gut microbial communities often lead to reduced diversity and altered metabolic outputs^8,9^. Among the key mediators of the host-microbiome crosstalk are bile acids (BAs), a class of amphipathic steroid molecules produced from cholesterol in the liver^10,11^. In humans, BA synthesis primarily proceeds via the classic pathway initiated by cholesterol 7α-hydroxylase (CYP7A1), yielding the host-derived bile acids (HBAs, also known as primary BAs) cholic acid (CA) and chenodeoxycholic acid (CDCA). Conjugation of HBAs to glycine or taurine in hepatocytes enhances their solubility and facilitates their function in lipid emulsification. Following their release into the small intestine during digestion, most BAs are reabsorbed in the terminal ileum and returned to the liver, completing a highly efficient recycling system that maintains BA homeostasis^12,13^.

Approximately five percent of BAs escape reabsorption and enters the colon^14^, where they are subjected to extensive microbial transformation. Gut bacteria encode a diverse array of enzymes that expand the chemical diversity and functional repertoire of BAs. Bile salt hydrolases (BSHs) deconjugate BAs, while enzymes encoded by the bile acid-inducible (*bai*) operon mediate 7-dehydroxylation to generate microbe-derived BAs (also known as secondary BAs) such as deoxycholic acid (DCA) and lithocholic acid (LCA). In parallel, hydroxysteroid dehydrogenases (HSDHs) catalyze stereo- and site-specific oxidation and reduction reactions, producing a wide range of epimerized BAs including ursodeoxycholic acid (UDCA), isoDCA and isoLCA^15^. Additional transformations include re-conjugation to various amino acids by BSH and dehydrogenation by site-specific reductases, which further increase the structural diversity of the BA pool^16,17^.

Microbial modification of BAs changes their chemical properties and consequentially, their signaling capabilities and function within the host. Microbe-derived BAs have been shown to regulate immune function, with epimerized forms such as isoLCA, isoDCA and UDCA promoting anti-inflammatory responses^18,19^. Global shifts in BA composition have been linked to intestinal inflammation and disease. In IBD, prior studies reported reduced levels of microbe-derived BAs alongside accumulation of host-derived precursors, suggesting impaired 7-dehydroxylation^4,20,21^. However, the extent to which additional microbial transformations contribute to IBD-associated BA pool remodeling and affect host physiology remains unclear.

Here, we performed a large-scale investigation of microbial BA metabolism in IBD by integrating publicly available metagenomic and metabolomic datasets alongside validation cohorts. We used AI-driven data mining for hypothesis generation combined with experimental validation in host and microbial systems to identify new mechanistic links between the gut microbiome and disease pathophysiology. By systematically characterizing functional microbial shifts in IBD, we found an unexpected role for HBA as drivers of dysbiosis independently of intestinal inflammation. We also uncovered a context-dependent role for microbial enzymes that mediate BA epimerization in producing dysbiosis-associated metabolites that disrupt host BA homeostasis.

## Results

### Bile acid epimerization is a hallmark of IBD

Intestinal inflammation is associated with alterations in both microbial community composition and BA profiles, particularly during active IBD^21,22^. To systematically investigate these changes at scale, we leveraged data from four published clinical studies that generated paired fecal shotgun metagenomic and metabolomic datasets: PROTECT^23^, PRISM^4^, iHMP^21^, and 1000IBD^9^. We curated a custom database comprising approximately 11,000 microbial enzymes involved in BA metabolism based on published literature^24–27^. After we quantified BA metabolism genes in metagenomic datasets, we trained neural networks to identify microbial gene modules associated with BA clusters (Extended Data Fig. 1a,b). Importantly, although our machine learning approach relied only on nominal BA identifiers, we uncovered four chemically coherent BA clusters: cluster a contained HBA with and without taurine/glycine conjugation; cluster b was marked by epimerized HBA and their microbe-derived conjugates; cluster c encompassed 7-dehydroxylated BAs, and cluster d included 7-dehydroxylated BAs that were re-conjugated by microbial enzymes (Fig. 1a). Among the models tested, the neural network trained on the PRISM dataset achieved the highest performance in predicting stool BA composition from fecal metagenomic profiles across cohorts (Extended Data Fig. 1b) and was therefore selected for downstream analyses. Genes in our custom BA metabolism database outperformed non-BA-related microbial genes in predicting stool BA composition and exhibited stronger prediction capabilities for BAs compared to other fecal metabolites (Extended Data Fig. 1c,d). Together, these findings indicate that our approach successfully captured the link between microbial BA enzymatic potential and fecal BA composition.

**Figure 1:**
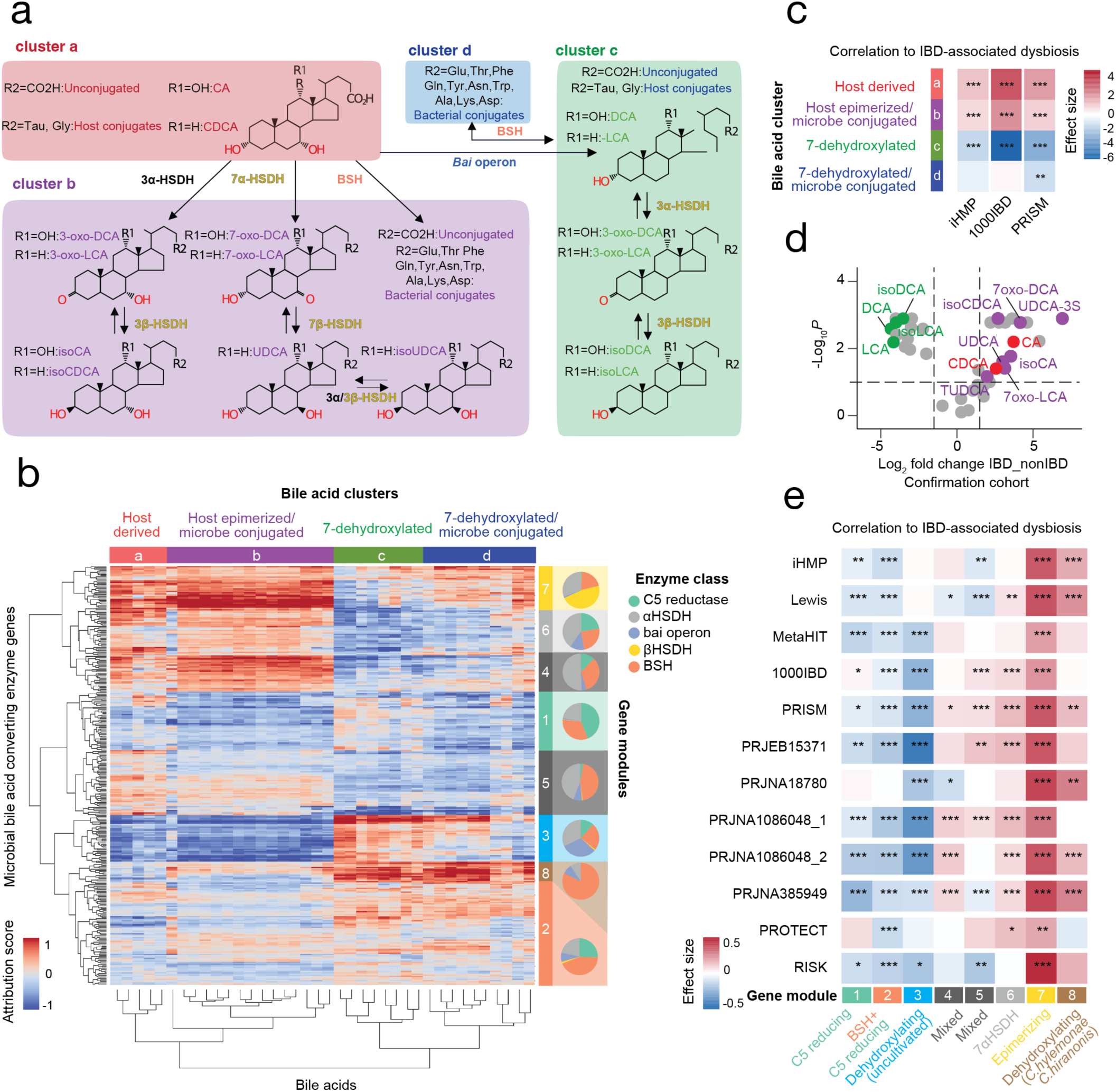
Epimerization of host derived bile acids is a hallmark of IBD-associated dysbiosis. (**a,b**) Stool metagenomic sequencing reads from four large inflammatory bowel disease (IBD) datasets (PRISM, iHMP, PROTECT and 1000IBD, total n= 1116) were mapped to a custom microbial bile acid (BA) converting enzyme gene database. Neural networks were trained to predict BA composition in paired metagenomics-metabolomics samples. **a**) Simplified schematic of bacterial BA metabolism based on BA clusters (clusters a-d) identified using neural networks. Enzyme classes: *BSH*, Bile salt hydrolase; *HSDH*, hydroxysteroid dehydrogenase; *Bai* operon, *Bai* genes A-I, responsible for C7 dehydroxylation. (**b**) Unbiased clustering of microbial BA converting enzyme genes (gene modules, row) and individual BAs (BA clusters, column) based on feature attribution scores inferred from the IBD dataset with the highest predictive power (PRISM). (**c**) Correlation between BA clusters and IBD-associated dysbiosis in indicated cohorts. (**d**) Targeted MS-based quantification of BAs in confirmation cohorts. (**e**) Correlation between BA epimerizing gene modules and IBD-associated dysbiosis across multiple studies. Statistical significance in (**c-e**) was determined with MaAslin2 based on the relative abundance of BA clusters (**c**), individual BAs (**d**) or mean clr-normalized gene module abundances (**e**). Sample size and epidemiological data of all cohorts in **Supplementary Table 1**. For further details on statistical analyses, see **Methods**.

To define functional operational units, microbial genes significantly predicting BA composition were grouped into BA metabolism gene modules based on their attribution scores (Fig. 1b). We further assigned BA predicting genes into enzyme classes based on their reported activity and structural conservation to better interpret the function of each gene module. Representative protein sequences were clustered at 85% amino-acid identity, yielding 4,866 representative proteins that were used for structural investigation with AlphaFold 3 (Extended Data Fig.1 e,f). BSH enzymes formed the largest number of structurally distinct subgroups, likely reflecting their broad phylogenetic distribution and varied substrate preference^28^. *bai F* enzymes which are part of the *bai* operon formed two closely related but distinct groups with segregated occurrence in either gene module 3 or 8 (Extended Data Fig.1g). Overall, BA metabolism enzymes within the same class showed high structural homology, validating our function-based grouping.

Finally, we observed that our model learned biologically meaningful associations between gene modules and BA clusters. For example, BA gene module 3, which showed the strongest contribution from *bai* genes, was positively associated with BA cluster c, encompassing 7-dehydroxylated BAs. In contrast, the HSDH-dominated gene module 7 was positively associated with cluster b, which contained epimerized HBAs (Fig. 1b). Gene module 7 was also associated with the HBA-rich cluster a, which includes the precursors of epimerized HBAs. Notably, gene modules positively associated with HBAs or epimerized HBAs were inversely correlated with clusters enriched in 7-dehydroxylated or microbe-conjugated 7-dehydroxylated BAs. These observations are consistent with the substrate product relationship among these BA groups and the role of microbial genes in mediating their interconversion.

Having validated our machine-learning based approach, we next examined the association between BA clusters and IBD-related dysbiosis. Samples were classified as either eubiotic or dysbiotic based on their median Bray–Curtis distance from the reference group within each study^21^ (Extended Data Fig. 2a,b). Consistent with previous reports^20,21^, IBD-related dysbiosis was characterized by a reduction in 7-dehydroxylated BA cluster c, accompanied by an enrichment in HBA cluster a (Fig. 1c). Notably, the epimerized HBA cluster b was also enriched in dysbiotic individuals, suggesting that HBAs and epimerized HBAs covary and that the remodeling of the intestinal BA pool in IBD extends beyond changes in the C7-dehydroxylation pathway. Leveraging the predictive power of our model, we then analyzed nine additional IBD datasets to assess the relationship between BA clusters and dysbiosis. Consistent with our findings in the matched metagenomics–metabolomics cohorts (Fig. 1c), clusters a and b were positively associated with dysbiosis across most studies examined, whereas cluster c showed a negative association (Extended Data Fig. 3a,b). Within each cluster, most individual BAs that could be consistently detected or predicted across all 12 studies were independently associated with IBD-related dysbiosis (Extended Data Fig. 3c). Because not all BAs with identical exact masses were resolved in our untargeted metabolomics training datasets, we were not able to unambiguously distinguish certain BA species such as UDCA and HDCA. To address this limitation and validate the enrichment of epimerized HBAs, we profiled stool BAs in an independent clinical IBD cohort (AUT, n = 40; cohort characteristics in Supplementary Table 1) using targeted LC–MS. HBAs including CA and CDCA as well as their epimerized derivatives isoCA, isoCDCA, UDCA and isoUDCA were enriched in stool samples from patients with IBD. In contrast, 7-dehydroxylated BAs including DCA, LCA and their isomerized forms such as isoDCA and isoLCA were depleted (Fig. 1d). These findings confirm that HBAs predominate over 7-dehydroxylated species in IBD and indicate that HBA epimerization is a pervasive feature of IBD-associated dysbiosis.

We next assessed whether the BA gene modules identified through our unsupervised approach were associated with IBD-related dysbiosis. Modules 6, 7, and 8 were positively associated with dysbiosis, with module 7 exhibiting the largest effect size and a significant association across all 12 studies examined (Fig. 1e and Extended Data Fig. 4a,b). Consistent with this observation, a random-effects meta-analysis (REML) identified a significant positive association between module 7 and IBD-related dysbiosis and diversity loss (Table 1). To further characterize alterations in microbial BA metabolism in IBD, we performed an in-silico perturbation analysis and evaluated the contribution of each BA metabolizing enzyme gene class to dysbiosis (Extended Data Fig. 4a). Within BA gene module 7, α- and β-HSDHs acting at the 3-OH and 7-OH positions, but not at the 12-OH position, contributed significantly to the positive association with dysbiosis (Extended Data Fig. 4c,d). We hereafter refer to module 7 as the BA-epimerizing gene module. Among the bacterial species encoding 3/7-HSDHs associated with IBD-related dysbiosis were *Escherichia coli* and *Mediterraneibacter gnavus* (formerly *Ruminococcus gnavus*) (Extended Data Fig. 4e), which were previously reported as enriched in IBD^29,30^. Because our primary analyses combined UC and CD, we next asked whether the association between module 7 and dysbiosis was driven by one IBD subtype. In a subtype-stratified sensitivity analysis, module 7 remained significantly associated with dysbiosis across both UC and CD cohorts, supporting a generalizable relationship between this gene module and IBD-related dysbiosis (Extended Data Fig. 5a,b). Collectively, these results suggest that dysbiosis in IBD and BAD is characterized by a loss of 7-dehydroxylation capacity, increased abundance of HBAs and epimerized HBAs, and an overrepresentation of microbial HSDHs (Extended Data Fig. 5c).

**Table 1:**
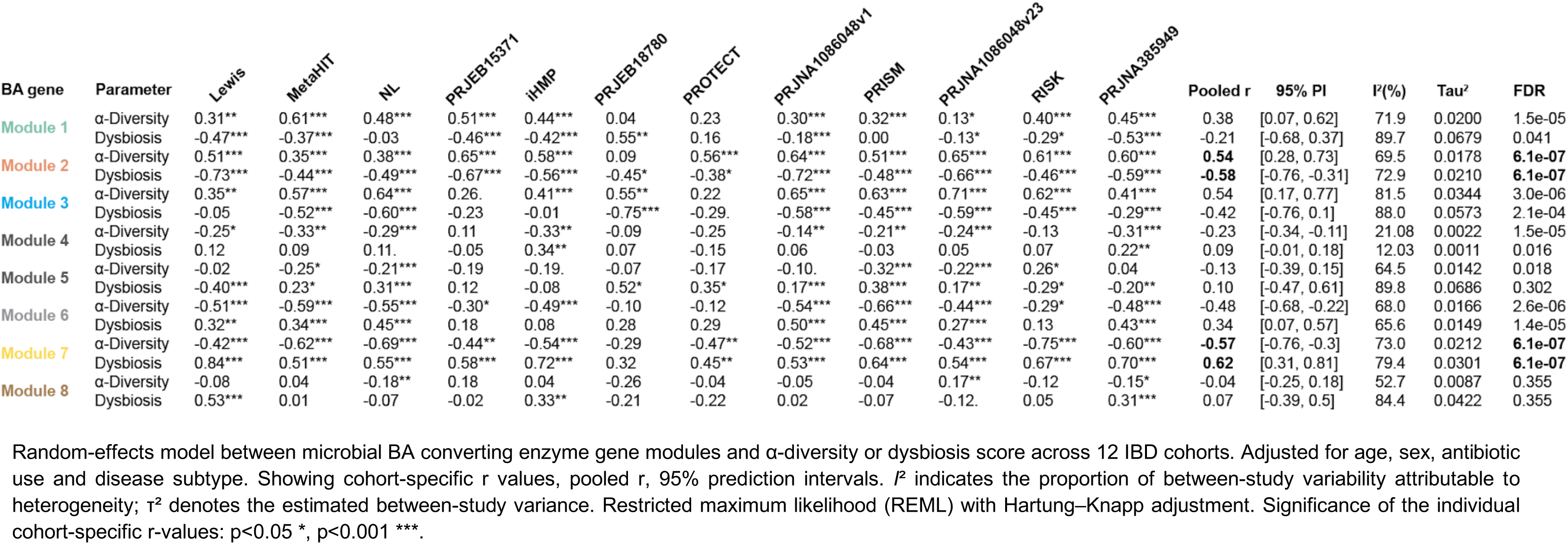
Random-effects meta-analysis of the correlation between IBD-associated dysbiosis / diversity loss and neural-network defined microbial BA converting gene modules.

### High levels of HBAs are associated with dysbiosis in the absence of overt inflammation

Since BA clusters were highly associated with dysbiosis scores (Table 2), we next tested whether they could explain additional variation in ecological microbiome metrics after adjusting for intestinal inflammation. We fitted linear models for dysbiosis score and Shannon diversity index including study, IBD-subtype, fecal calprotectin levels and BA cluster composition. Together, these features explained approximately 60% of sample variance (R²) in dysbiosis scores and Shannon diversity among IBD patients. Compared to fecal calprotectin, BA composition alone explained a large fraction of the variance (Fig. 2a and b, left). Consistently, a meta-analysis showed that adding BA composition to models already containing fecal calprotectin and IBD subtype (ΔR²) increased the explained variance in individual studies by approximately 30 percent on average (Fig. 2a and b, middle). In secondary analyses including only individual BA clusters, HBA cluster a and 7-dehydroxylated BA cluster c showed the largest incremental ΔR² values (Fig. 2a and b, right). Since the relative abundance of these clusters represent two mutually exclusive BA pool configurations either dominated by precursors (cluster a) or products (cluster c), these analyses suggest that increased relative abundance of HBAs is an independent contributor to dysbiosis and loss of microbial diversity.

**Figure 2:**
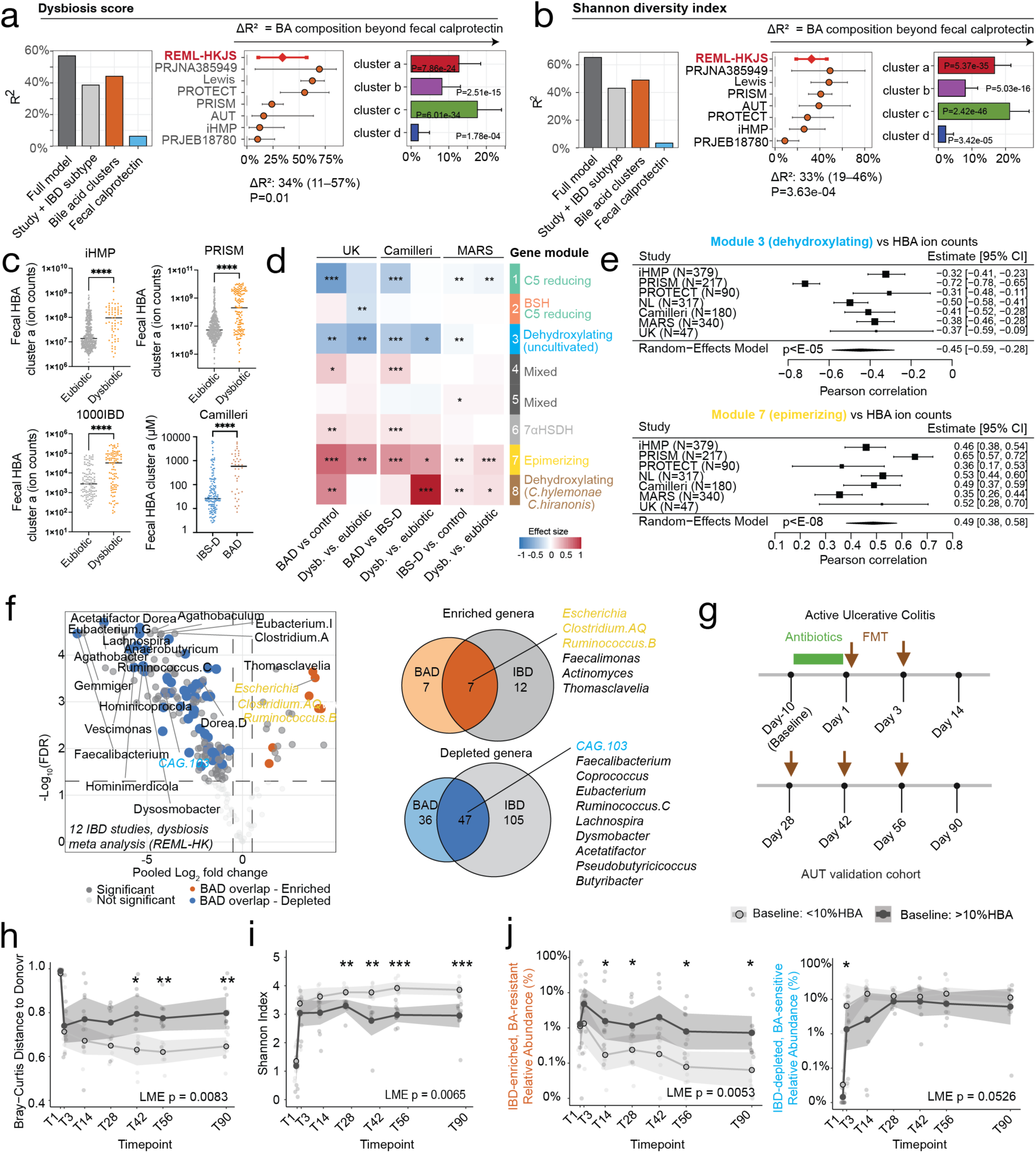
Increased host derived BAs explain dysbiosis and microbial diversity loss in inflammatory bowel disease. Impact of bile acids (BA) on dysbiosis score (**a**) and Shannon diversity index (**b**) variance in IBD cohorts with accompanying fecal calprotectin data (PRJNA385949, Lewis, PROTECT, PRISM, iHMP, PRJEB18780, AUT confirmation cohort). (**a,b**) Left: variance (R²) explained by linear models based on study + IBD subtype (baseline), BA composition, fecal calprotectin or the factors combined (baseline + BA composition + fecal calprotectin combined=full). Middle: random-effects meta-analysis across cohorts of the incremental variance explained by BA composition beyond fecal calprotectin (ΔR² = R²_full_− R²_fCal+base_). Right: incremental variance explained by individual BA clusters beyond fecal calprotectin. (**c**) Fecal host-derived BA levels (HBA, cluster a) in IBD and diarrhea-predominant irritable bowel syndrome (IBS-D) with or without bile acid diarrhea (BAD). Showing within study comparisons of IBD-associated dysbiosis versus eubiotic controls (iHMP, PRISM and1000IBD) and IBS-D with or without BAD (Camilleri). (**d**) Differential abundance of neural-network defined BA converting gene modules in BAD (UK + Camilleri cohorts) and IBS-D (Mars cohort). (**e**) Meta analysis of correlation between HBA levels and microbial BA converting enzyme module 3 (BA dehydroxylating-uncultivated, top) or module 7 (BA epimerizing, bottom) across IBD and BAD/IBS-D studies. (**f**) Intersection of IBD- and BAD-associated dysbiosis. Volcano plot showing bacterial genus level random-effects meta-analysis across 12 IBD cohorts comparing dysbiotic vs. eubiotic subjects. Showing pooled log_2_ fold-change (x-axis) and log_10_ FDR (y-axis) (middle). Bacterial genera overlapping with BAD in the same direction are highlighted in color. Venn diagrams show the number of taxa depleted (left, blue) or enriched (right, orange) in IBD-related dysbiosis and BAD.(**g**) Study design of the AUT confirmation cohort. Patients with active UC received 10 days of antibiotics followed by five ileoscopic fecal microbiota transplants (FMT) and were subjected to longitudinal stool sampling at the indicated time points. (**h**) Bray Curtis distance to donor over time after FMT, stratified for baseline (day -10) signs of BA malabsorption (>10% relative abundance of HBA in dark grey, <10% HBA in light grey). Lower distance indicates greater FMT engraftment. (**i**) Shannon diversity trajectories after FMT stratified by baseline BA status. (**j**) Longitudinal trajectories of IBD-enriched bile acid-resistant genera (left) and IBD-depleted bile acid-sensitive genera (right) after FMT. Statistical significance was determined via linear models (**a,b,e,h,i,j**), a Restricted maximum likelihood (REML) meta-analysis with Hartung–Knapp adjustment (**a,b,e**), a Mann Whitney-U test (**c,h,i,j**), MaAslin2 (**d,f**) or a Pearson-correlation test (**e**). Sample size and epidemiological data of all cohorts in **Supplementary Table 1**. For further details on statistical analyses, see **Methods**.

**Table 2:**
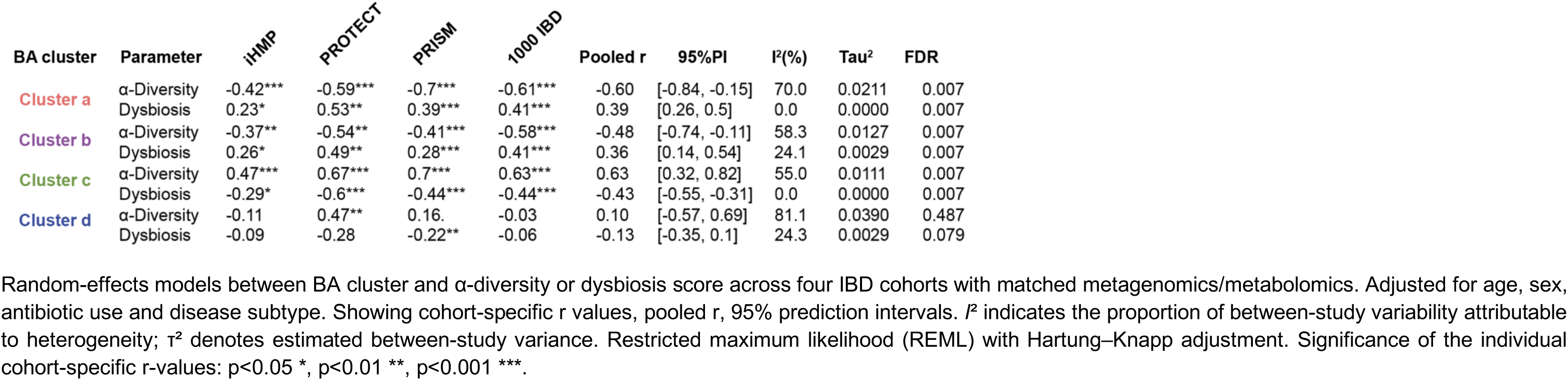
Random-effects meta-analysis of the correlation between IBD-associated dysbiosis / diversity loss and neural-network defined BA clusters.

**Table 3:** Random-effects meta-analysis of UC disease activity / inflammation and neural-network defined HBA cluster a and epimerized HBA cluster b.

| BA cluster | Parameter | iHMP | PROTECT | PRJNA385949 | PRISM | Pooled r | 95% PI | I<sup>2</sup>(%) | Tau<sup>2</sup> | FDR |
| --- | --- | --- | --- | --- | --- | --- | --- | --- | --- | --- |
| Cluster a | Calprotectin | 0.36 | 0.36* | 0.45 | 0.19 | 0.29 | [0.12, 0.44] | 0 | 0 | 0.033 |
|  | Disease Activity | 0.34 | 0.19 | 0.43 | - | 0.26 | [0.02, 0.48] | 0 | 0 | 0.056 |
| Cluster b | Calprotectin | 0.34 | 0.32 | 0.78 | 0.02 | 0.21 | [-0.24, 0.59] | 14.7 | 0.0091 | 0.143 |
|  | Disease Activity | 0.21 | 0.17 | 0.37 | - | 0.19 | [0.08, 0.29] | 0 | 0 | 0.033 |

Random-effects models between HBA cluster a and epimerized HBA cluster b versus fecal calprotectin and disease activity (Mayo disease activity score) across the indicated UC cohorts. Restricted maximum likelihood (REML) with Hartung–Knapp adjustment. Corrected for age, sex and antibiotic use. Showing cohort-specific r values, pooled r, 95% prediction intervals. I<sup>2</sup> indicates the proportion of between-study variability attributable to heterogeneity; $\tau^2$ denotes estimated between-study variance. Significance of the individual cohort-specific r-values: \*p<0.05. BA composition was predicted from metagenomics in the PRJNA385949 cohort.

**Table 4:** Random-effects meta-analysis of CD disease activity / inflammation and neural-network defined HBA cluster a and epimerized HBA cluster b.

| BA cluster | Parameter | Lewis | iHMP | PRJNA385949 | PRISM | PRISM | Pooled r | 95% PI | I<sup>2</sup>(%) | Tau<sup>2</sup> | FDR |
| --- | --- | --- | --- | --- | --- | --- | --- | --- | --- | --- | --- |
| Cluster a | Calprotectin | 0.14 | -0.13 | 0.45 | -0.09 | 0.45 | 0.04 | [-0.25, 0.32] | 11.4 | 0.0046 | 0.909 |
|  | Disease Activity | 0.1 | 0.02 | -0.13 | - | - | 0.06 | [-0.08, 0.2] | 0 | 0 | 0.501 |
| Cluster b | Calprotectin | 0 | -0.16 | 0.42 | -0.14 | 0.02 | -0.07 | [-0.2, 0.07] | 0 | 0 | 0.501 |
|  | Disease Activity | -0.01 | 0.04 | -0.25 | - | - | 0 | [-0.15, 0.15] | 0 | 0 | 0.975 |

Random-effects models between HBA cluster a, epimerized HBA cluster b and fecal calprotectin or disease activity (Crohn's disease activity index) across the indicated CD cohorts. Restricted maximum likelihood (REML) with Hartung–Knapp adjustment. Corrected for age, sex and antibiotic use. Showing cohort-specific r values, pooled r, 95% prediction intervals. I<sup>2</sup> indicates the proportion of between-study variability attributable to heterogeneity; $\tau^2$ denotes estimated between-study variance. Significance of the individual cohort-specific r-values: \*p<0.05. BA composition was predicted from metagenomics in the Lewis and PRJNA385949 cohort.

Next, we leveraged additional patient cohorts to dissect putative causal relationships between high HBA levels and dysbiosis. Idiopathic BA diarrhea (BAD) is a functional GI disorder characterized by elevated fecal HBA levels (*i.e.*, >10% of total stool BAs^31^), increased stool frequency and no overt inflammatory pathology. Similarly to BAD patients, individuals with IBD-related dysbiosis showed increased fecal HBA concentrations relative to eubiotic controls (Fig. 2c). We also observed similarities between IBD-related dysbiosis and BAD at the metagenomics level. The BA epimerizing gene module 7 was increased in two independent BAD cohorts relative to their respective controls, while the BA dehydroxylating gene module 3 was reduced. Importantly, these changes were not soley driven by the presence of diarrhea as the microbiome of patients with BAD still displayed a significant enrichment in HBAs and the BA epimerizing module when compared to those from patients with diarrhea-predominant irritable bowel syndrome (IBS-D, Camilleri cohort^32^) (Fig. 2d). In a random effect model-controlled metanalysis, HBA concentration was negatively correlated with the abundance of the dehydroxylating BA gene module 3 and positively correlated with the epimerizing module 7 across multiple IBD and BAD cohorts (Fig. 2e), suggesting that the link between HBA and changes in microbial BA transformation is not contingent on intestinal inflammation. Given the observed similarities between IBD and BAD, we then set out to identify microbes commonly affected by high HBA levels by performing a taxonomical meta-analysis on the 12 IBD datasets and intersecting bacterial genera differentially abundant in BAD and healthy controls (UK cohort^33^). Approximately 1/3 of the taxa depleted or enriched in IBD were similarly affected in BAD (Fig. 2f). Due to lower microbial diversity in both clinical conditions, we found a greater number of shared depleted microbes (47 genera) relative to enriched (7 genera). Shared depleted taxa included the uncultivated BA-dehydroxylating CAG-103 and short-chain fatty acid producing bacteria such as *Faecalibacterium*, *Butyribacter* and *Lachnospira*. Amongst the bacteria enriched in both IBD and BAD we identified *E. coli* and group B *Ruminococcus* (which includes *M. gnavus*), major contributors to the BA epimerizing gene module 7. Altogether, these analyses indicate that high HBA levels are associated with changes in composition and function of the microbiome regardless of the presence of intestinal inflammation.

### HBA drive intestinal dysbiosis and select for bacteria carrying HSDHs

The overlap between the intestinal microbiome signatures seen in both IBD and BAD suggested that high levels of HBAs could be an inflammation-independent factor shaping microbial ecology in these patients. To explore this idea, we carried out longitudinal microbiome analyses in a subset of patients in our IBD confirmation cohort undergoing fecal microbiota transfer (FMT) for the treatment of active UC (Fig. 2g). Despite similar clinical parameters (Supplementary Table 1), patients presenting high levels of HBA (>10%) at baseline (10 days before FMT) showed significantly lower engraftment as measured by the Bray-Curtis distance to graft and had reduced microbial diversity post-transplant relative to patients with lower HBA levels (Fig. 2h,i). Further, we observed an enrichment in BA-resistant bacteria and a transient depletion of BA-sensitive species (defined in Fig.2f) in patients with high baseline HBA levels (Fig. 2j), suggesting that high HBAs in IBD can push eubiotic microbial communities towards dysbiotic states. We next asked whether BA-targeted interventions could reverse IBD/BAD-associated microbiome changes. For this, we leveraged longitudinal data from a BAD cohort undergoing treatment with the BA-sequestrant Colesevelam (Extended Data Fig. 5d). As expected, BAD patients had increased levels of BA-resistant bacteria and reduced abundance of BA-sensitive taxa at baseline and showed a significant increase in the later at 4 weeks post treatment (Extended Data Fig. 5e,f). These results further support the notion that high levels of HBAs are sufficient to drive part of the dysbiotic signatures seen in IBD/BAD.

To directly test this hypothesis, we cultivated healthy donor stool samples in mini bioreactor arrays (MBRAs)^34^ in the presence of an intermediate (0.5mM) or high (5mM) concentration of CA+CDCA (Fig. 3a), corresponding to the range found in the ileum of dysbiotic / BAD individuals (Fig. 2c)^35,36^. This continuous flow culture model allows for daily longitudinal sampling over multiple time points (T, see Experimental set up in Fig. 3a) including during stabilization (baseline, T0+T2), exposure to BAs (T3-T6), and washout (T8+T10). HBAs changed microbial community structures compared to baseline in a dose- and time-dependent manner (Fig. 3b). High levels of HBAs had a strong effect on microbial diversity (Fig. 3c), depleting BA-sensitive commensals and enriching for bacteria overrepresented in IBD/BAD (Fig.2f) such as *E. coli* and *M. gnavus* (Fig. 3d). Although CAG-103 was not detected at baseline, high HBA levels led to a loss of other BA-sensitive commensals with positive effects on human health such as *Faecalibacterium* and *Butyribacter* (Fig. 3e).

**Figure 3:**
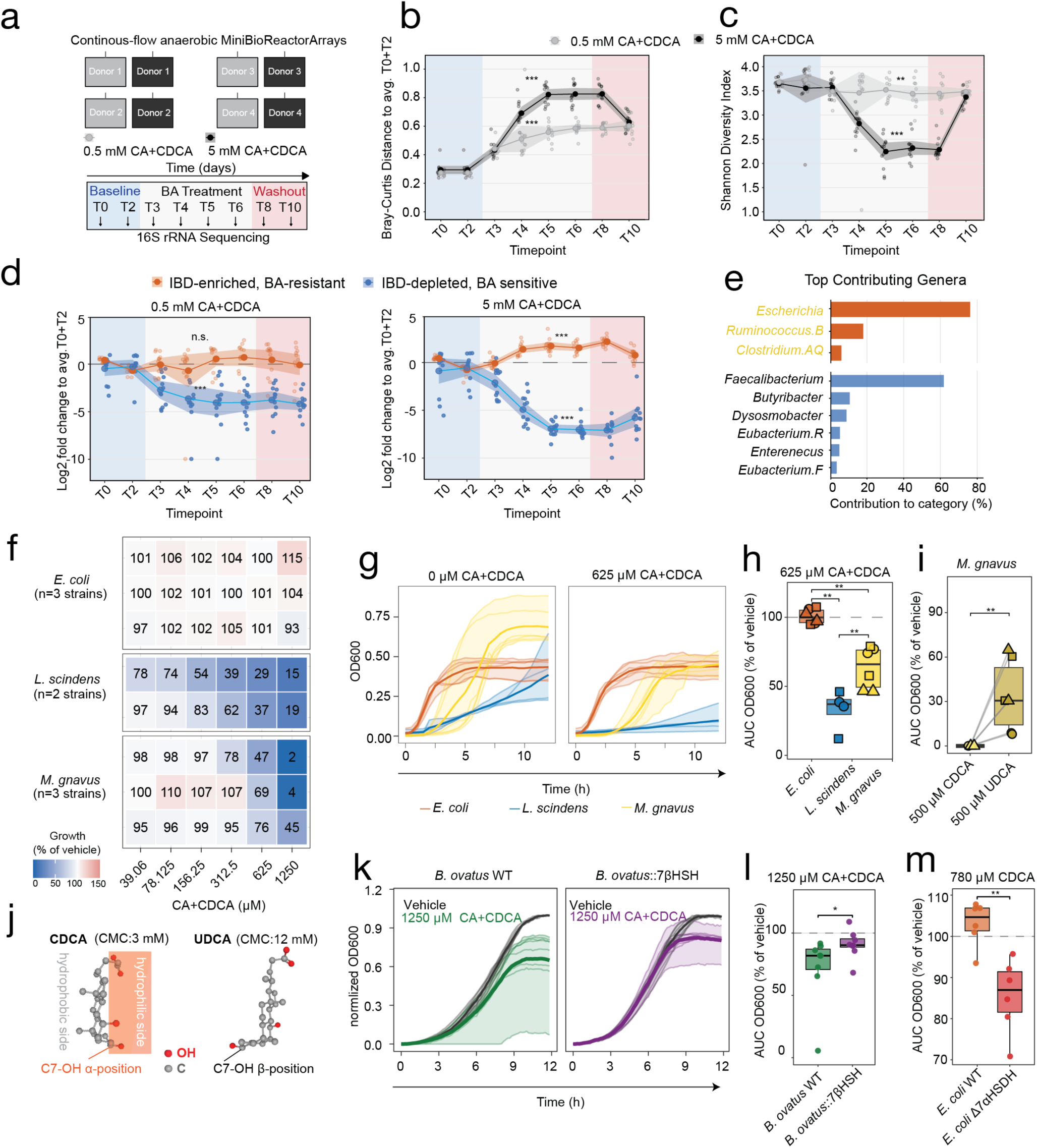
Host derived BA exposure recapitulates IBD-associated diversity loss and bloom of BA epimerizing strains. (**a**) Experimental schematics for data shown in (**b-e**): four healthy donor stool samples were inoculated in triplicates in continuous flow anaerobic MiniBioReactor Arrays (MBRA). After a two-day stabilization period, MBRA were treated for four days with high (5 mM, dark grey) or intermediate (0.5 mM, light grey) concentrations of HBAs (CA+CDCA, 2:1 ratio) and bacterial communities were assessed by 16S RNA sequencing. Bray-Curtis distance to each reactor’s baseline composition (avg. T0+T2, **b**) and Shannon diversity index (**c**) upon HBA exposure. (**d-e**) HBAs select for BA-resistant (orange) and deplete BA-sensitive (blue) bacterial genera shared between BAD/IBD-dysbiosis. Showing log_2_ fold-change from baseline (avg. T0+T2) on the y-axis and time point on the x-axis. (**e**) Top genera contributing to the BA-resistant and BA-sensitive signatures during the treatment period. Bacteria highlighted in yellow carry hydroxysteroid dehydrogenases (HSDHs) found in the BA-epimerizing gene module 7. (**f**) Heatmap of *L. scindens, M. gnavus* and *E. coli* strains growth rates under a gradient of host-derived BAs (CA+CDCA, 2:1 ratio). Showing area under the curve (AUC) of OD600 over 12 hours normalized to vehicle. (**g,h**) Strain-specific growth in the presence of 1250 µM CA + CDCA. Thick lines in (**g**) denote average growth of strains of indicated species. Symbol shapes in (**h**) denote different strains. (**i**). *M. gnavus* growth at 500 µM of CDCA versus UDCA. Showing OD600 AUC relative to the vehicle control. (**j**) PubChem 3D structures of CDCA (<u>CID:10133</u>) and UDCA (<u>CID:31401</u>) showing a break in the hydrophilic (alpha) and hydrophobic (beta) sides of CDCA upon C7 hydroxyl-group epimerization. CMC, critical micelle concentration. (**k,l**) Growth of an engineered *Bacteroides ovatus* strain carrying 7βHSDH from *M. gnavus* (*B. ovatus::7βHDSH*) and wild type (WT) parental control strain under 1250 µM CA + CDCA. Showing growth curves (**k**) and OD600 AUC relative to vehicle (**l**). (**m**) Growth of WT and 7αHSDH-deficient (*hdhA::Kn^r^*) *E.coli* under 780µM CDCA. Showing OD600 AUC relative to the vehicle control. To determine statistical significance in (**b-d**), data from each BA concentration were analyzed separately using linear mixed-effects models with phase (*Baseline, Treatment, Washout*) as a fixed effect and reactor as a random intercept. Displaying P values corresponding to Bonferroni adjusted Baseline versus Treatment contrast. Data shown in (**f-m**) are representative from two independent experiments. (**f-h, j**) n=6 for *E. coli*, n=4 for *L. scindens* and n=6 for *M. gnavus*. (**k-m**) n=8 biological replicates. Significance was determined with a Wilcoxon rank-sum (**h**) or two-sided Mann-Whitney U test (**j,l,m**). *p<0.05, **p<0.01, ***p<0.001.

Next, we assessed microbe-intrinsic responses to HBAs by measuring the growth of multiple *E. coli*, *M. gnavus*, and *Lachnoclostridium scindens* isolates across a range of CA and CDCA concentrations (Fig. 3f). Because CAG-103 remains uncultivated, we used two *L. scindens* strains as representatives of 7-dehydroxylating bacteria. *L. scindens* was the most sensitive to HBAs, exhibiting marked growth impairment even at low concentrations. *M. gnavus* displayed intermediate tolerance, whereas *E. coli* remained resistant at all concentrations tested, including the highest concentration used in this assay (1.25 mM).

Previous studies have shown that hydroxysteroid dehydrogenases (HSDHs) promote bacterial survival in the presence of 7-dehydroxylated BAs by converting them into epimers with reduced detergent activity. We therefore hypothesized that HSDHs encoded by *E. coli* and *M. gnavus* contribute to their expansion under high HBA conditions. Consistent with this hypothesis, exposure to UDCA had a substantially weaker inhibitory effect on *M. gnavus* growth relative to CDCA (Fig. 3i-j)^37^. Moreover, the inhibitory activity of BA decreased with increasing hydroxyl-group epimerization, following the order CDCA > isoCDCA > UDCA > isoUDCA (Extended Data Fig. 6a). Because genetic manipulation of *M. gnavus* remains challenging, we used *Bacteroides ovatus* as a chassis to directly test whether HSDHs enhance bacterial fitness in the presence of CDCA. The genome of *B. ovatus* encodes a 7α-HSDH. Complementation of the 7-HSDH pathway by knock-in of the *M. gnavus* 7β-HSDH enabled conversion of CDCA to UDCA by *B. ovatus* and significantly improved growth in the presence of CDCA (Extended Data Fig. 6b,c and Fig. 3k,l). Similarly, *B. ovatus* expressing an active 3α/β-HSDH module from *M. gnavus*^18^ exhibited enhanced growth in CDCA compared to the parental WT strain (Extended Data Fig. 6d,e). Finally, although *E. coli* was generally resistant to elevated HBA concentrations, deletion of its 7α-HSDH abolished HBA C7 oxidation and resulted in a significant growth defect in the presence of CDCA (Extended Data Fig. 6b,c and Fig. 3m). Together, these findings indicate that HSDHs enhance bacterial growth under high HBA conditions and provide a mechanistic explanation for the enrichment of *E. coli* and *M. gnavus* observed in IBD/BAD.

### Increased HBA levels are associated with colonic epithelial remodeling in UC

Having established that elevated HBA levels contribute to dysbiosis in IBD, we next investigated whether 7-hydroxylated BAs were also associated with other disease features. Crohn’s disease (CD) and ulcerative colitis (UC) differ markedly in their clinical manifestations and tissue involvement, with the first being typified by transmural, discontinuous lesions throughout the gastrointestinal tract and the latter being characterized by relatively superficial and continuous colonic inflammation. While disease activity measurements in UC include an objective endoscopic assessment (Mayo score), in CD this is restricted to patient-reported parameters, further limiting our ability to directly compare these two settings^38,39^. We thus carried out separate REML meta-analyses to identify potential associations between BA clusters and disease parameters in the two IBD subsets. In patients with UC, we found that HBA cluster a correlated positively with calprotectin and disease activity and that the epimerized HBA cluster b was associated with disease activity (Tables 3). No significant associations were identified in CD patients (Tables 4).

To further investigate the relationship between BAs and host pathophysiology in UC, we analyzed single-cell RNA-sequencing data from colonic biopsies obtained from in a subset of our validation cohort (reported elsewhere; manuscript in revision). Within this sub cohort, HBA cluster a was associated with UC disease activity (Extended Data Fig. 7a), indicating that the relationships observed in larger public UC cohorts were preserved in this group.

Recovered intestinal cell populations were broadly categorized into epithelial (secretory and absorptive), immune (T cells, myeloid cells, and B/plasma cells), and stromal/vascular (endothelial, stromal, and fibroblast) compartments based on transcriptional clustering (Fig. 4a). We then asked whether HBA cluster a and the epimerized HBA cluster b were associated with cellular composition after accounting for mucosal inflammation. To address this, we quantified incremental ΔR² gained by adding individual BA cluster clr values to an endoscopic Mayo score baseline model in a hierarchical composition screen. Variance in patient-level cell type composition within the epithelial compartment, specifically within the secretory lineage, was associated with cluster a abundance (Fig. 4b, Extended Data Fig. 7b, left). Within the secretory epithelial compartment, goblet cell (GC) frequencies were positively correlated with cluster a abundance at the patient level(Extended Data Fig. 7b, right). We also validated this association between HBA levels and GC abundance in an independent UC cohort using histological analysis of Periodic Acid–Schiff (PAS)-stained tissue (Fig. 4c). Increased numbers of GCs further correlated with less voluminous mucus granules per cell (Fig. 4d), suggesting potential changes in GC maturation state.

**Figure 4:**
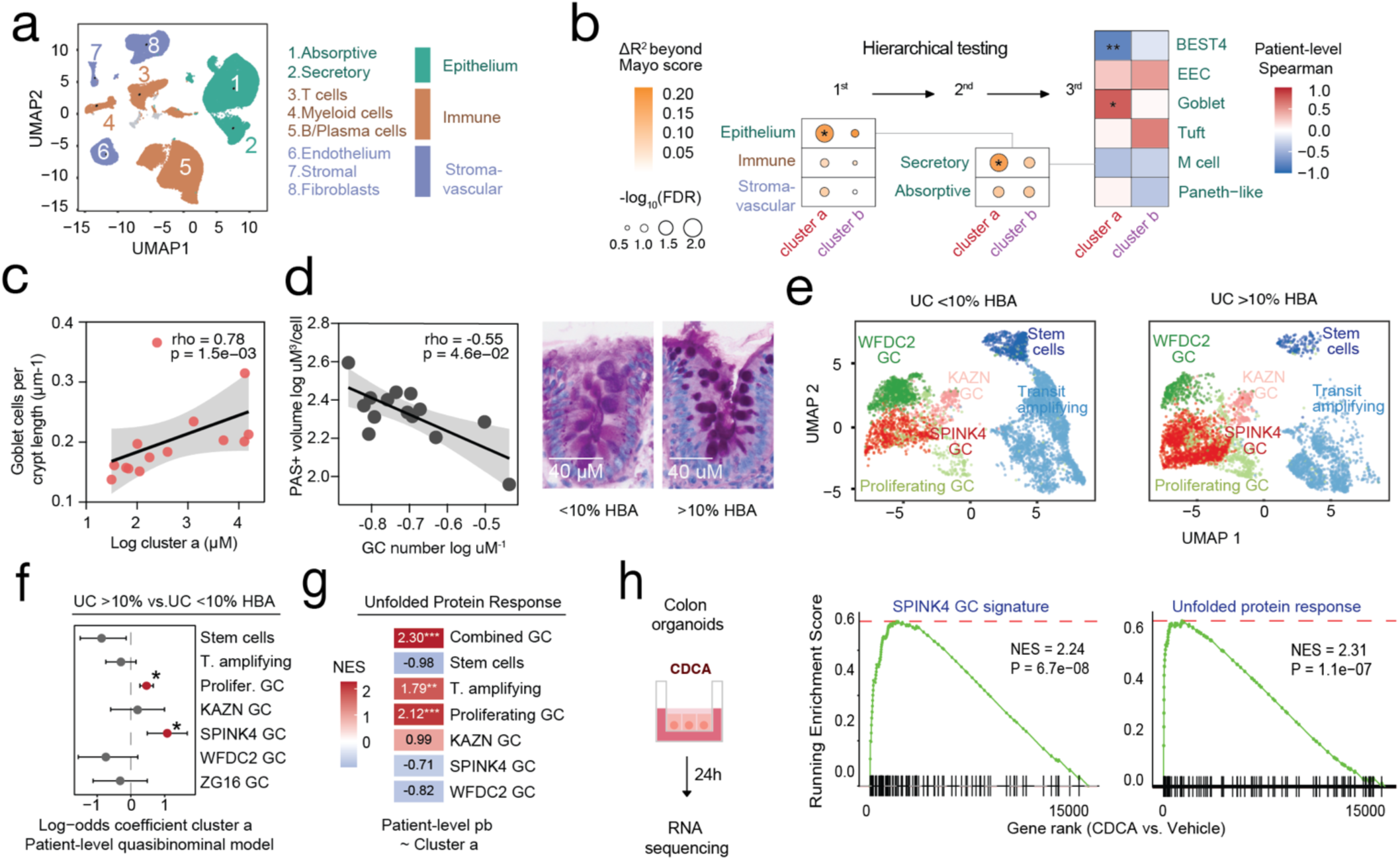
Host derived bile acids link dysbiosis to goblet cell remodeling and epithelial stress in ulcerative colitis. (**a**) Uniform Manifold Projection and Approximation (UMAP) representation of single-cell RNA sequencing profiling of UC patient’s colonic biopsies, colored by major cellular compartments. (**b**) Hierarchical assessment of cell type composition changes associated with BA pool remodeling. Cell type abundance within each level was modeled as isometric log-ratio (ILR)-transformed compositions and tested against the indicated fecal BA cluster abundances with Mayo score as a covariate. Bubble color indicates incremental ΔR^2^ beyond endoscopic Mayo score; bubble size indicates FDR. Heatmap showing patient-level Spearman correlations between the indicated secretory epithelial cell subsets and BA clusters. (**c,d**) Independent histological validation of the BA-goblet cell interaction in UC. (**c)** Correlation between fecal HBA cluster a abundance and goblet cell density normalized to crypt length. (**d)** Left: Correlation between goblet cell density normalized to crypt length and mucus load assessed by PAS-positive area within individual cells. Right: representative colonic PAS-stained sections from UC patients with high (>10%) or low (<10%) fecal HBA cluster a abundance. (**e**) UMAP of stem, transit-amplifying, and goblet cell maturation states in UC patients with HBA cluster a >10% versus <10% of the total fecal BA pool. **(f**) Patient-level correlation of HBA cluster a with goblet cell maturation states. Positive coefficients indicate enrichment with higher HBA cluster a abundance. (**g**) Gene expression changes across the goblet cell maturation stages. Patient-level pseudobulk profiles were generated for combined and individual maturation states, and genes were ranked by association with HBA cluster a abundance (Extended Data Fig.7). Showing Reactome Unfolded Protein Response pathway normalized enrichment scores (NES) across the indicated epithelial cell populations. (**h**) Organoid validation of SPINK4-like goblet-state program and BA-induced stress response. Paired patient-derived colon organoids were treated with either CDCA or vehicle and subjected to bulk RNA-sequencing. Genes were ranked by differential expression (CDCA vs. vehicle) and tested by GSEA. Showing enrichment for the GOBLET_SPINK4 signature defined from goblet cell maturation states shown in (**e,f**) and for the Reactome UPR pathway. For clustering parameters used to generate the UMAP shown in (**a**), see **Methods.** (**a,b,e-g**) n=38 colonic biopsies from 10 patients spanning different endoscopic Mayo scores. (**c,d**) n=14 UC patients with matched colonic biopsies and stool BA profiles from an independent cohort, statistical significance via spearman correlation. (**h**) n=3 independent biological replicates. In (**b**) cell-composition outcomes were represented by ILR-transformed within-branch composition coordinates, and reported R^2^ values summarize variance explained across ILR axes. (**b,f-h**) FDR values were calculated using Benjamini-Hochberg correction. *p<0.05, ***p<0.001.

To investigate how high HBAs impacted GC differentiation in UC, we visualized their maturation states in patients with high (>10%) versus low (<10%) HBA levels and tested for associations with HBA cluster a (Fig. 4e,f). Higher abundance of cluster a was significantly associated with a shift towards proliferative and SPINK4^+^ GC states, with the latter being previously implicated in mucosal regeneration during chronic intestinal inflammation. An endoscopic Mayo-adjusted biopsy-level random-effects model showed concordant effects, with nominal significance for proliferating and SPINK4^+^ GC (Extended Data Fig. 7c). Next, we performed Reactome pathway gene set enrichment analysis (GSEA) on patient-level pseudo bulk gene expression across GC differentiation states. In addition to interferon signaling, pathways involved in GC maturation and mucus secretion including vesicular trafficking, glycosylation and the unfolded protein response (UPR) were significantly associated with HBA cluster a abundance (Extended Data Fig. 7d). UPR-associated gene expression in transit-amplifying cells and proliferating GCs was also associated with HBA cluster a abundance at the patient level (Fig. 4g). To determine whether HBAs are sufficient to induce these pathways in immature epithelial cells, we treated healthy tissue derived colonic organoids with CDCA and performed RNA sequencing. CDCA treatment upregulated pathways related to endoplasmic reticulum stress and the UPR (Fig. 4h and Extended Data Fig. 7e), suggesting that HBAs are sufficient to induce part of the transcriptional programs observed in HBA-high UC patients. Although mature GC depletion has been previously reported as part of the epithelial remodeling during UC^40,41^, these findings suggest that within affected patients elevated HBA levels increase the abundance of immature and proliferating GC by promoting stress responses.

### Epimerized HBAs disrupt FXR signaling and the BA negative feedback loop

In ileal enterocytes, Farnesoid X receptor (FXR) activation by agonistic BAs induces the expression of fibroblast growth factor 19 (FGF19), which is then released into the portal circulation. In hepatocytes, FGF19 activates FGFR4 to suppress CYP7A1, thereby maintaining BA homeostasis through a negative feedback mechanism (Fig. 5a). The elevated levels of HBAs observed in IBD-related dysbiosis suggested disruption of this regulatory axis. To investigate how altered microbial BA transformation may affect host BA metabolism, we leveraged ileal gene expression datasets with paired fecal BA profiles from the iHMP study. Patients with CD often present ileitis or undergo small intestine surgical resection^42^, which can directly affect BA absorption; therefore, we restricted our analyses to UC cases. We generated a FXR activation gene expression signature by treating healthy tissue ileal organoids with the FXR agonist GW4064 (Fig. 5b). GSEA was then used to compare FXR pathway activity in ileal biopsies from UC patients with low (<10%) versus high (>10%) fecal HBA levels. GW4064-induced genes (FXR-up signature) were significantly downregulated in patients with high HBA levels, whereas genes repressed by GW4064 (FXR-down signature) were upregulated, consistent with reduced FXR activity in this group (Fig. 5c). Further, while healthy individuals showed a positive correlation between ileal FXR activation signature and fecal CDCA levels (Fig. 5d, left), this relationship was absent in patients with UC (Fig. 5d, right), suggesting that despite their increased abundance in UC, HBA acids fail to effectively induce FXR signaling in this setting.

**Figure 5:**
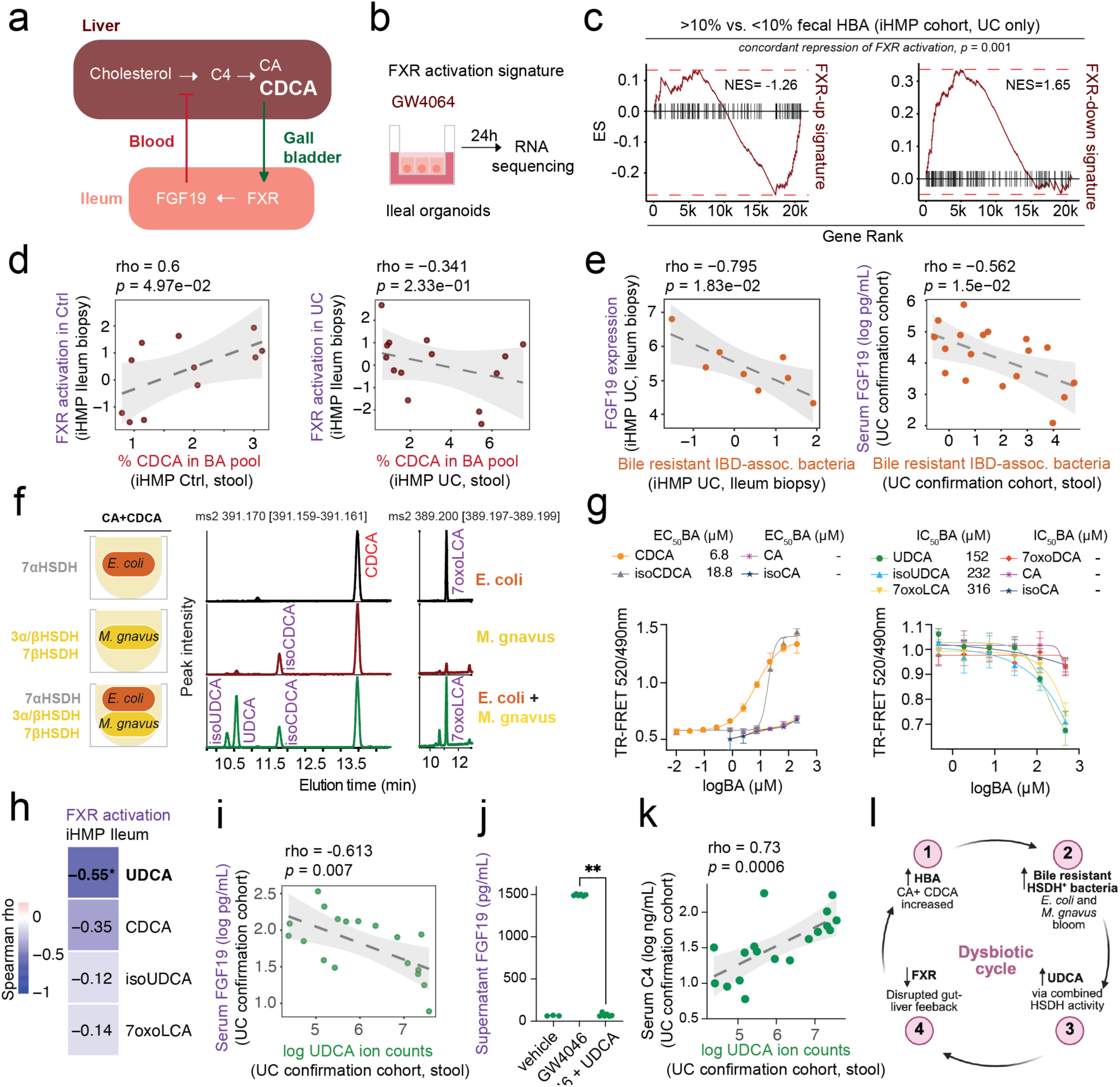
Dysbiosis-associated epimerized host derived BAs impair the FXR–FGF19 axis in ulcerative colitis. (**a**) Simplified depiction of the enterohepatic BA feedback loop: host derived bile acids (HBA) are synthesized in the liver from cholesterol with 7α-hydroxy-4-cholesten-3-one (C4) as intermediate. After secretion from the gall bladder into the gastrointestinal tract, HBAs are actively taken up by ileocytes and activate the Farnesoid X Receptor (FXR) in the nucleus. FXR activation induces FGF19 secretion into the portal blood; FGF19 signaling in the liver downregulates HBA synthesis. (**b**) Generation of an FXR activation gene expression signature. Ileal organoids were treated with the FXR agonist GW4064 at 10µM for 24h and subjected to RNA-sequencing. (**c**) GSEA analyses for the GW4064-upregulated (induced by FXR activation) and -downregulated (repressed by FXR activation) gene signatures amongst the differentially expressed genes in ileal biopsies from UC patients (iHMP cohort) with high (>10%) vs. low (<10%) fecal HBA levels. (**d**) Correlation between fecal CDCA abundance (x-axis) and ileal FXR activation (y-axis) inferred from FXR gene signature enrichment in healthy individuals (left) or patients with UC (right). (**e**) Correlation between FGF19 levels (x-axis) and BA-resistant IBD-associated bacteria (y-axis) in patients with UC from the iHMP (left) and validation cohorts (right). (**f**) *E. coli* and *M. gnavus* were grown alone or in combination in the presence of HBA (CA+CDCA 100µM) overnight. BA transformation in the supernatants was assessed by LC/MS. Showing mass spectrometry traces for each condition and the peaks for the indicated BAs at their characteristic retention time. (**g**) FXR co-activator recruitment assay with recombinant FXR ligand binding domain (LBD). Showing the time-resolved fluorescence resonance energy transfer (TR-FRET, y-axis) upon incubation with BAs at indicated concentrations (x-axis). Left, agonist mode; right, antagonist mode (+ 10µM CDCA). (**h**) Correlation between selected fecal BA levels and ileal FXR activation in patients with UC (iHMP dataset). (**i**) Correlation between fecal UDCA (x-axis) and serum FGF19 (y-axis) levels in the UC confirmation cohort. (**j**) Ileal organoid monolayers were treated with 0.6µM GW4064 with or without 1000µM UDCA and FGF19 production was measured by ELISA. (**k**) Correlation between fecal UDCA (x-axis) and serum C4 (y-axis) levels in the UC confirmation cohort. (**l**) Schematic of proposed “dysbiotic cycle”: increased HBAs select for BA-resistant bacteria such as *E. coli* and *M. gnavus*. These HSDH-carrying bacteria produce UDCA that disrupts BA enterohepatic negative feedback loop, further enhancing HBA synthesis. For details on RNA-sequencing analysis shown in **b** and **c** see **Methods**. Statistical significance in **d, e, h,** and **k** was determined with a Spearman Correlation test or two-sided Mann-Whitney U test (**j**). (**c, d, e** left and **h)** n=14 UC patients and 11 control subjects from the iHMP cohort with matching ileum biopsy 16S, ileum bulk RNA-sequencing and stool BA data. (**e**,right, **i** and **k**) n= 18 UC patients from confirmation cohort.

Notably, ileal FGF19 expression showed a strong negative correlation with the enrichment of BA-resistant, IBD-associated bacterial taxa as defined by our taxonomic IBD/BAD meta-analysis (Fig. 2f and Fig.5e) implying that these microbes may impair FXR signaling in the intestine via BA transformation. To explore this hypothesis, we cultured *E. coli* and *M. gnavus* individually or together in the presence of CA and CDCA and profiled the resulting BA composition. *M. gnavus* monocultures produced isoCA and isoCDCA, consistent with its expression of 3α/3β-HSDH (Fig. 5f). *E. coli* alone generated 7-keto derivatives of CA and CDCA (7-keto DCA and 7-keto LCA, respectively), whereas co-culture of *E. coli* and *M. gnavus* produced a broader array of modified BAs, including isoCA, isoCDCA, UDCA, and isoUDCA. Importantly, co-culture derived BAs were strongly enriched in IBD validation cohorts (Fig. 1d), suggesting that these microbes contribute to the BA shifts observed in IBD- and BAD-associated dysbiosis.

We next tested whether microbially modified BAs influence FXR signaling using a FRET-based FXR co-activator recruitment assay. As expected, CDCA was a substantially more potent FXR agonist than CA (Fig. 5g, left). In contrast, isoCDCA exhibited a nearly threefold higher EC_50_ than CDCA (18.8 μM vs. 6.8 μM), indicating that epimerization of the 3-OH group reduced FXR activation. In antagonist mode, UDCA, isoUDCA, and 7-oxo-LCA significantly reduced CDCA-induced FXR activation (Fig. 5g, right), with IC_50_ values of 152 μM, 232 μM, and 316 μM, respectively. FXR activation in ileal tissue from UC patients was strongly negatively correlated with fecal UDCA levels (Fig. 5h), consistent with its antagonistic activity reported in human studies^43^. Moreover, fecal UDCA abundance was inversely correlated with circulating FGF19 in the validation cohort (Fig. 5i), suggesting impaired FXR-driven feedback signaling under high UDCA conditions. To directly test this notion, we measured FGF19 secretion in small intestinal organoids treated with GW4064 at EC_80_. UDCA significantly reduced GW4064-induced FGF19 secretion compared with vehicle (Fig. 5j), indicating that epimerized HBAs can directly disrupt the FXR–FGF19 axis.

Circulating levels of the BA precursor 7α-hydroxy-4-cholesten-3-one (C4), an intermediate in the hepatic BA synthesis pathway, are increased in IBD^44,45^. We found that serum C4 levels were positively correlated with fecal UDCA abundance (Fig. 5k), implying that high levels of this microbe-derived BA derepress hepatic BA synthesis in patients with UC. Together, these findings suggest that BA epimerization by BA-resistant pathobionts such as *E. coli* and *M. gnavus* attenuates FXR signaling and disrupts the intestinal–hepatic negative feedback loop, thereby sustaining a high-HBA state (Fig. 5l).

## Discussion

Decreased microbial diversity and shifts in immunomodulatory microbial metabolites including BA have been consistently reported in various IBD studies^22^. However, despite several efforts, the underlaying mechanisms and ecological drivers of these changes remain poorly understood. Conventional statistics can be used to validate differential abundances of individual genes or metabolites but often fail to capture higher-order interactions, non-linear dependencies and coordinated, many-to-many relationships^46,47^. This is especially relevant for BAs, where one metabolite can reflect the combined activity of sequential bio-transformations carried out by multiple bacterial taxa. Here, we applied a deep learning framework specifically designed to extract latent structures from high-dimensional microbiome data. In combination with our comprehensive microbial BA transforming enzyme database, this strategy allowed us to identify microbial BA-converting enzyme modules linked to biologically meaningful BA clusters. Our analyses confirmed previous studies showing that C7-dehydroxylated BAs are depleted in IBD and linked this deficit to a reduced abundance of *bai* genes encoded by uncultivated Firmicutes^20,26^. Consistent with earlier reports, we also observed an enrichment of HBAs in IBD-associated dysbiosis^4,19^. Importantly, we identified a previously unrecognized class of microbially modified BAs associated with IBD: epimerized HBAs. While these metabolites were nearly undetectable in healthy controls, epimerized HBAs including UDCA were consistently enriched in IBD-associated dysbiosis across three independent public datasets and our validation cohort. Moreover, we found that a BA-epimerizing gene module enriched in HSDHs from pathobiont-associated species such *as E. col*^48^, *C. innocuum*^49^, and *M. gnavus*^29^. This module was consistently associated with both epimerized HBA levels and IBD-related dysbiosis across 12 independent datasets. Together, these findings indicate that BA remodeling in IBD is characterized not only by the loss of C7-dehydroxylation capacity but also by a coordinated gain of BA epimerization activity, leading to the accumulation of epimerized HBAs as a hallmark metabolic feature of IBD-associated dysbiosis.

The structural heterogeneity of BA-metabolizing enzymes, epitomized by bile salt hydrolases, highlights the evolutionary diversity of microbial BA metabolism and suggests that convergent metabolic functions can be encoded by phylogenetically distinct taxa. Interestingly, structurally distinct *bai* genes from *Clostridium hiranonis* and *Clostridium hylemonae* were associated with dysbiotic microbiome states in some of the analyzed studies. Nevertheless, our findings indicate that the dominant BA-dehydroxylating bacteria supporting gastrointestinal homeostasis are likely represented by yet uncultivated taxa. Expanding culturomics approaches to recover these organisms will be critical for elucidating their biology and evaluating their therapeutic potential.

Using harmonized data and the predictive capabilities of our neural network models we could perform a stringent meta-analysis on microbial dysbiosis/diversity loss, fecal BA composition and GI inflammation in IBD. We found that BA cluster explain a substantial proportion of variance in microbial dysbiosis score and diversity independently of inflammatory status. The presence of key features of IBD-associated dysbiosis in the non-inflammatory condition idiopathic BAD supports a role for elevated HBAs as ecological filters that drive microbial community changes independently of inflammation. In both settings, elevated HBAs were linked to an enrichment of the BA-epimerizing gene module and expansion of *E. coli* and *M. gnavus*, while commensals such as SCFA producers and uncultivated *bai*-carrying Firmicutes were depleted. Additionally, we found that IBD patients with high HBA levels showed lower engraftment and reduced microbial diversity upon fecal microbiota transplantation. Conversely, patients with BAD showed signs of microbiome recovery upon Colesevelam treatment, suggesting that targeted approaches might improve dysbiosis in clinical settings of excess HBA. Together, these findings support the notion that HBA excess is not only a disease biomarker but also a direct driver of IBD-related dysbiosis.

Previous studies reported that HBAs promote *Enterobacteriaceae* growth by modulating host factors^50^. Here, we show that HBAs also directly act on the microbiome. Exposure of complex microbial communities to disease-relevant concentrations of CA and CDCA induced dose-dependent reductions in diversity and selected for BA-resistant organisms that are overrepresented in IBD and BAD. Putative SCFA-producing BA-sensitive bacterial taxa such as *Lachnospira* and *Faecalibacterium* were depleted, whereas HSDH-containing bacteria such as *E. coli* and *M.gnavus* bloomed. By testing pure cultures of several type strains and clinical isolates, we found that *E.coli* and *M. gnavus* are intrinsically more tolerant to HBAs, while BA-dehydroxylating species are relatively sensitive. Using engineered *B. ovatus* strains we show that HSDH enzymes contribute to bacterial fitness under BA stress, likely by epimerizing toxic HBA into less detergent forms. *E.coli* and *M. gnavus* were sufficient to convert CDCA into the epimerized HBA forms we found enriched in IBD (*i.e.,* 7oxoLCA, UDCA, isoUDCA, isoCDCA). Taken together, our findings provide a molecular explanation for IBD-related dysbiosis and the associated changes in BA profile.

Patients with CD who undergo ileal resection frequently display elevated fecal HBA levels^42^. However, the increase in HBAs observed in UC and BAD is less readily explained and has been suggested to arise from microbiome-dependent mechanisms^51^. Unexpectedly, our meta-analysis found that elevated fecal HBA levels correlated with disease activity in UC but not CD. Since fecal calprotectin is a less reliable marker for intestinal inflammation in CD then UC, it remains possible that HBA affect CD pathophysiology and this association was not captured in our analyses. Colonic single-cell transcriptomic and histological analyses revealed associations between HBA levels and secretory epithelial remodeling in UC, including goblet cell and tuft cell programs, that persisted after accounting for endoscopic inflammation and microbiota composition. These findings suggest that HBAs may contribute to epithelial remodeling through mechanisms that are at least partly independent of inflammatory burden.

Beyond their ecological effects on the gut microbiome, our findings suggest that epimerized HBAs directly influence host signaling pathways in the ileum. Although HSDH-carrying bacteria including *M. gnavus* and *E.coli* were enriched in ileal biopsies of patients with high fecal HBA in published cohorts, we have not directly assessed this in our confirmation groups. Nonetheless, microbe-derived BA are absorbed systemically, undergo hepatic conjugation and become part of the enterohepatic circulation pool, suggesting that these molecules could interfere with FXR activation in the small intestine^52,53^.

We discovered that the positive relationship between fecal HBA levels and ileal FXR activation observed in healthy individuals was lost, indicating impaired gut–liver feedback regulation. HSDH-encoding, BA-resistant bacteria such as *E. coli* and *M. gnavus* were associated with reduced ileal FXR signaling, decreased FGF19 expression, and increased BA synthesis as measured by serum C4. Mechanistically, we found the that epimerized HBAs UDCA and isoUDCA act as FXR antagonists. These results support a model in which microbial BA epimerization attenuates FXR signaling and FGF19 secretion, thereby promoting BA overproduction and establishing a self-reinforcing cycle of elevated HBA levels and dysbiosis.

In summary, our study identifies excess fecal HBAs as a major ecological driver of IBD-associated dysbiosis. We demonstrated that elevated HBAs deplete BA-sensitive commensals and select for BA-resistant HSDH-encoding pathobionts that disrupt host BA feedback signaling. Our findings have direct translational relevance, as approved BA sequestrants could potentially be repurposed to lower luminal BA stress in the colonic lumen and precondition patients for microbiome-restorative therapies.

## Acknowledgments

We thank Joris van der Veeken, Abigail E Rose, Christoph Bock, Stephan Reichel and all the members of the Campbell and Bergthaler lab for helpful discussions. We thank Evette B M Hillman, Julian Walters, Paula J Carlson and Michael Camilleri for providing raw data from their bile acid diarrhea cohorts. The Vienna BioCenter Core Facilities (VBCF) Metabolomics Facility acknowledges funding from the Austrian Federal Ministry of Education, Science & Research; and the City of Vienna. Research in the Campbell laboratory is supported by the Austrian Science Fund (FWF; 10.55776/COE7, 10.55776/FG29 and 10.55776/F7000), 2023 Federation of European Biochemical Societies (FEBS) Excellence Award, and the European Research Council (ERC) under the Horizon 2020 research and innovation program (ERC-2023-STG 101117175). Gregor Gorkiewicz acknowledges funding from the Austrian Science Fund (FWF) [doi.org/10.55776/COE7]. Benoit Chassaing’s laboratory was supported by a Starting Grant (grant agreement Invaders No. ERC-2018-StG-804135) and a Consolidator Grant (grant agreement InterBiome No. ERC-2024-CoG-101170920) from the European Research Council (ERC) under the European Union’s Horizon 2020 research and innovation program, ANR grants EMULBIONT (ANR-21-CE15-0042-01) and DREAM (ANR-20-PAMR-0002), grant from the AFA Crohn RCH France and from the French government through the France 2030 investment plan managed by the National Research Agency (ANR), as part of the ANR-24-PESA-0008 and ANR-23-IAHU-0012.

## Author contributions

CC and MB conceptualized the study and wrote the original draft of the manuscript. CC supervised the project and acquired funding. AF, PS, SL, CH, CG, CP, WR, MT, CG, GB, and GG recruited patients and curated clinical cohorts. AM and CG isolated patient-derived bacterial strains from colonic biofilms. FS, TKN, TA, FP, EK, and MB performed single-strain bacterial in vitro experiments. MG, PH, GB, CF, and MB conducted organoid in vitro experiments. MD, BC, and MB performed bacterial MBRA experiments and analyzed 16S rRNA amplicon sequencing data. MB carried out data curation as well as metagenomic and transcriptomic bioinformatic analyses. GB and MB performed single-cell RNA sequencing analyses. ML and GW conducted protein structure prediction and analysis. AC, TK, and MB performed bile acid analyses. All authors corrected and approved the final version of this manuscript.

## Methods

### Bile acid converting enzyme database and metagenomics analysis

We built a comprehensive database of bacterial bile acid converting enzyme genes starting from published datasets including, bile salt hydrolases (BSH)^24^, hydroxysteroid dehydrogenases (HSDH)^25^, bai-genes^20,26^ and C5 reductases^27^. Additionally, experimentally validated genes^25,27^ were used for NCBI blastp searches against nr_cluster_seq with the following parameters: 50% identity, 80% coverage for bai-genes and C5 reductases, according to *Jin WB et al.* 60% identity and 80% coverage was used to distinguish different HSDH subtypes (i.e. 7aHSDH, 3aHSDH)^25^. Furthermore, hmm models for HSDH subtypes and C5 reductases were created to screen the Human Reference Gut Microbiome v2^54^ protein catalog using hmmer. The resulting protein sequences were clustered at 85% identity and shortBRED-identify^55^ with uniref90 (v2025-04) background was used to create unique markers. To validate our database we predicted the structure of 4866 enzymes with Alphafold3^56^. The FASTA sequence of each enzyme was used to generate a multiple sequence alignment. We generated 5 diffusion samples per enzyme using a randomly chosen seed, selecting the structure with the highest ‘ranking_score’ for downstream analysis. To focus our analysis on high confidence predictions, we filtered the structures on Alphafold3 confidence metrics: mean plddt (predicted local distance difference test) larger than 75, mean pae (pairwise alignment error) smaller than 10 Å and pTM (predicted template modeling score) larger than 0.75 and the ranking_score (aggregate score from multiple confidence metrics) larger than 0.8 (**Extended Data Fig. 1e**). Pairwise structural similarities were computed for all-against-all structures using Foldseek^57^ in TM-align mode. Hierarchical clustering on the resulting TM-score distance matrix (1-TM-score) using the computed alnTM-score and the Ward method implemented in SciPy confirmed enzyme subclass specific conservation in our final database (**Extended Data Fig. 1f**). The optimal number of BSH structural clusters was determined to be five using the maximum silhouette criterion.

To harmonize published metagenomics studies we created a HPC job array pipeline to process SRA, consisting of: quality control with fastp (paired-end processing, duplicate removal, read length threshold 60 bp), host-contaminants removal with Bowtie2 (sensitive mode with hg37_dev as reference), MetaPhlAn 4.2.2^58^ (database: mpa_vJan25_CHOCOPhlAnSGB_202503) for taxonomic profiling and ShortBRED-quantify^55^ for BA converting enzyme detection. The pipeline and BA database are publicly available at: **github.com/MaximilianBaumgartner/MetaBile**.

### Microbial bile acid converting enzyme neural network analysis

The MiMeNet^46^ package was adapted to run on NVIDIA H100-GPU infrastructure and used to predict bile acid composition from paired metagenomics ShortBRED output of four large IBD studies^4,9,21,23^. MiMeNet was run with standard settings: clr-transformation of metabolite and gene tables, 10 iterations of 10-fold cross-validation (100 individual predictions) and 100 iterations with 10-fold cross-validation (1000 individual predictions) for randomly shuffled background generation. Cross-dataset predictions were performed using the build-in “-external_micro and -external_metab” function. To compare MiMeNet performance of non-BA metabolizing genes 50 random prevalence and mean-detected-abundance percentile-matched subsets were drawn from a HUMAnN 3^59^ gene families table, filtered for our BA converting enzyme database via DIAMOND^60^ (pident >= 40, query coverage >= 0.70, subject coverage >= 0.50). The forked MiMeNet package, including models to predict BA from our BA converting enzyme database shortBRED results are publicly available at: **github.com/MaximilianBaumgartner/MiMeNet_tf2.** Sample size, read depth and epidemiological data of all cohorts are in **Supplementary Table 1**. Details on how many total enzymes per enzyme subclass are in the BA converting enzyme database, the number deemed significant to predict BA composition by our MiMeNet analysis, together with average RPKM values can be found in **Supplementary Table 2.** Individual BA, their chemical structure, in how many studies they were reported, relative abundance within the whole BA pool as well as neural-network prediction spearman correlation coefficients within and across studies can be found in **Supplementary Table 3.**

### Differential abundance analysis

We defined IBD associated “dysbiosis” as >90th median percentile taxonomic Bray-Curtis dissimilarity distance to controls^21^ (**Extended Data Fig.S2**). For differential abundance testing of BA cluster relative abundance or mean clr-normalized gene module abundances MaAslin2^61^ was used, in longitudinal datasets, “subject” was included as a random effect.

### In-silico perturbation analysis

Baseline associations between ML-defined module scores and dysbiosis status were estimated within each of the 12 IBD cohorts using MaAsLin2 as described above. To assess the contribution of individual enzyme classes, genes annotated as BSH, C5 reductases, *bai* genes, α-HSDHs, or β-HSDHs were computationally removed one class at a time. The remaining enzyme count matrix was re-normalized using CLR transformation and compositionally corrected to the unperturbed CLR space, after which module scores and module–dysbiosis associations were recalculated. Perturbation effects were defined as the change in log2 effect size relative to the unperturbed model. Study-level baseline and perturbation effects were combined across cohorts using random-effects meta-analysis with REML, and p-values were adjusted across tested module–enzyme class combinations using the Benjamini–Hochberg FDR procedure.

### IBD dysbiosis meta-analysis

Within each cohort, we calculated cohort-specific partial correlations between BA modules or BA clusters and Shannon diversity index, dysbiosis score, fecal calprotectin, and disease activity, adjusting for age, sex, and antibiotic exposure; combined UC+CD analyses were additionally adjusted for IBD subtype. As not all cohorts with clinical metadata had publicly available metabolomics, BA composition was predicted from metagenomics in the Lewis, PRJNA385949 and PRJEB18780 cohorts (see **Supplementary Table 1**). Cohort-specific correlation coefficients were then pooled using random-effects meta-analysis with restricted maximum likelihood (REML) and Hartung–Knapp adjustment. Between-study heterogeneity was summarized using *I*² and τ². Multiple testing was controlled using the Benjamini–Hochberg false discovery rate (FDR). To evaluate whether BA composition explained microbiome variation beyond intestinal inflammation, we modeled the four BA clusters as compositional data. Cluster abundances were first converted to proportions and transformed into isometric log-ratio (ILR) coordinates, yielding three orthogonal balances: (cluster b + cluster a) versus (cluster c + cluster d), cluster a versus cluster b, and cluster c versus cluster d. For dysbiosis score and Shannon diversity, we fit linear models including: IBD-subtype and study, fecal calprotectin alone, BA cluster composition alone, and a full model with all parameters. The additional variance explained by BA composition beyond fecal calprotectin was quantified as ΔR² = R²_full_ − R²_fCal_. Cohort-specific ΔR² estimates were derived by bootstrap resampling and pooled across cohorts using random-effects REML meta-analysis with Hartung–Knapp adjustment. As a secondary analysis, individual BA clusters were assessed separately. Each CLR-transformed cluster was added individually to models containing fecal calprotectin, and the corresponding incremental variance explained was reported.

### Meta-analysis of microbial taxa in IBD/BAD

MetaPhlAn SGBs were converted to GTDB genus level using the sgb_to_gtdb_profile.py. MaAslin2 was used on log-transformed relative abundance of GTDB genera to detect dysbiosis association in 12 large IBD cohorts. Cohort-specific genus-level effect estimates were extracted as log_2_ fold changes with corresponding standard errors and pooled using random-effects meta-analysis with REML and Hartung–Knapp adjustment. Genera were retained if detected in at least 50% of studies contributing to a given comparison. Significant genera were defined as those with FDR < 0.05, absolute pooled log_2_ fold change > 0.5, and support from at least three studies. To identify bile acid–linked microbial signatures, pooled IBD and dysbiosis results were compared with an independent bile acid diarrhea dataset. Detailed results from our taxonomic meta-analysis can be found in **Supplementary Table 4**.

### MBRA analysis

Fresh stools were collected at home in sealed anaerobic containers, transported on ice, processed within 2 h in an anaerobic chamber, aliquoted, and stored at −80 °C. The MBRA system was operated as previously described^34,62^, using 24 anaerobic, continuously stirred 15-mL reactors containing basal reactor medium (BRM), modified by the addition of 0.25% porcine mucin. After sterilization and 72 h anaerobic equilibration, reactors were inoculated with 3.8 mL of fecal slurry prepared from stools resuspended at 2.5% w/v in anaerobic PBS, vortexed, centrifuged, and filtered through a 100 µm mesh. Communities were equilibrated for 20 h before continuous flow was initiated at 1.875 mL/h, corresponding to an 8-h retention time; 400 µL samples were collected over time and stored at −80 °C. For each donor, two bile acid conditions were tested in BRM containing 0.25% porcine mucin: 0.5 M or 5 mM total host-derived bile acids, consisting of an equimolar 2:1 mixture of NaCA and NaCDCA. Filter-sterilized bile acid stocks were added post-autoclaving. After a 2-day stabilization, bile acid-supplemented medium was perfused for 5 days, followed by a 4-day washout period. Bacterial DNA was extracted from 50 µL MBRA suspension using the DNeasy 96 PowerSoil Pro QIAcube HT kit (Qiagen), with mechanical disruption via bead beating (TissueLyzer II, Qiagen). Microbial community composition was profiled by 16S rRNA gene amplicon sequencing of the V4 region using barcoded 515F/806R primers with Illumina adapters and Golay error-correcting sample barcodes (515F 5’-AATGATACGGCGACCACCGAGATCTACACGCTXXXXXXXXXXXXTATGGTAA TTGTGTGYCAGCMGCCGCGGTAA-3’ and 806R 5’-CAAGCAGAAGACGGCATACGAGATAGTCAGCCAGCCGGACTACNVGGGTWTCTAAT-3’).

PCR was performed with AccuStart™ II PCR SuperMix using 0.2 µM primers and 10–100 ng template DNA, with cycling conditions of 95 °C for 3 min; 30 cycles of 95 °C for 45 s, 50 °C for 60 s, and 72 °C for 90 s. Amplicons were purified with AMPure magnetic beads, and sequenced at the Genom’IC facility, Institut Cochin, Paris, France utilizing Illumina MiSeq 250PE (Illumina). Sequence demultiplexing, quality filtering and taxonomic classification were performed using the DADA2^63^ and the SBDI Sativa curated 16S GTDB database^64^. Microbiome shifts were analyzed using R, the phyloseq, microbiome and tidyverse package. Shannon diversity indexand Bray-Curtis distance was calculated on total sum scaled ASV-tables. For further analysis counts were agglomerated at the genus level and classified as “IBD-enriched, BA-resistant” or “IBD-depleted, BA-sensitive” based on our IBD taxonomic meta-analysis, genera present in <10% of samples were excluded. Timepoints were grouped into baseline, BA treatment, and washout phases, with reactors assigned to mid- or high-concentration bile acid conditions. Treatment effects were tested separately for each bile acid concentration and genus category using linear mixed-effects models with phase as a fixed effect and reactor as a random intercept, followed by estimated marginal means and Bonferroni-adjusted pairwise contrasts.

### Bacterial single strain growth curves

Bacterial strains were grown anaerobically overnight in BHIS and diluted to OD600 = 0.05. Bile acids were prepared as 2× BHIS stocks and twofold serially diluted in 96-well plates before addition of an equal volume of culture, yielding a starting OD600 of 0.025. Medium-only and blank controls were included, plates were sealed with gas-permeable membranes (BreathEasy), and cultures were incubated anaerobically at 37 °C with OD600 measured every 30 min for 12 h. Experiments included independent biological replicates from separate overnight cultures and/or colonies. Two E. coli strains were isolated from endoscopic colon brushes of healthy subjects. Additional reference strains were ordered from the German Collection of Microorganisms and Cell Cultures (DSMZ) or ATCC. Keio collection *E. coli hdhA::Kn^r^* and and the corresponding parental strain were ordered from Horizondiscovery as part of the Keio collection.

To complement the 7a-HSDH precent in *B. ovatus* and enable UDCA synthesis from CDCA, the 7b-HSDH (uniprotid:A7B4V1) genes Mg_7b were synthesized (IDT DNA) after codon-optimization for expression in *B. ovatus*. Golden Gate assembly kit (NEB) was used to assemble the following fragments in 5’ to 3’ order: ppWW3806, ribosome binding site #8 (RBS), Mg_7b open reading frame and pWW3810. The assembled construct was transformed into competent DATC *E. coli*^65^ and positive clones were selected and verified by whole-plasmid sequencing. The plasmid was conjugated into *B. ovatus* by mixing 1 mL each of overnight DATC donor (grown in LB + 0.3 mM 2,6-diaminopimelic acid) and *B. ovatus* (in BHI) recipient cultures, pelleting at 8,000 × g for 1 min, and resuspending the combined pellet in ∼100 µL residual medium. The suspension was spotted onto nonselective BHI + 0.3 mM 2,6-diaminopimelic acid agar without streaking, allowed to dry briefly, and incubated aerobically overnight at 37 °C, agar side down. Lawn was then scraped on a BHI double selection plate containing both gentamycin (200μg/mL) and erythromycin (25μg/mL), followed by anaerobic culture for 2 days at 37°C. Colonies were replated and genomic integration confirmed via whole genome sequencing.

The following strains and plasmids have been used in this study:

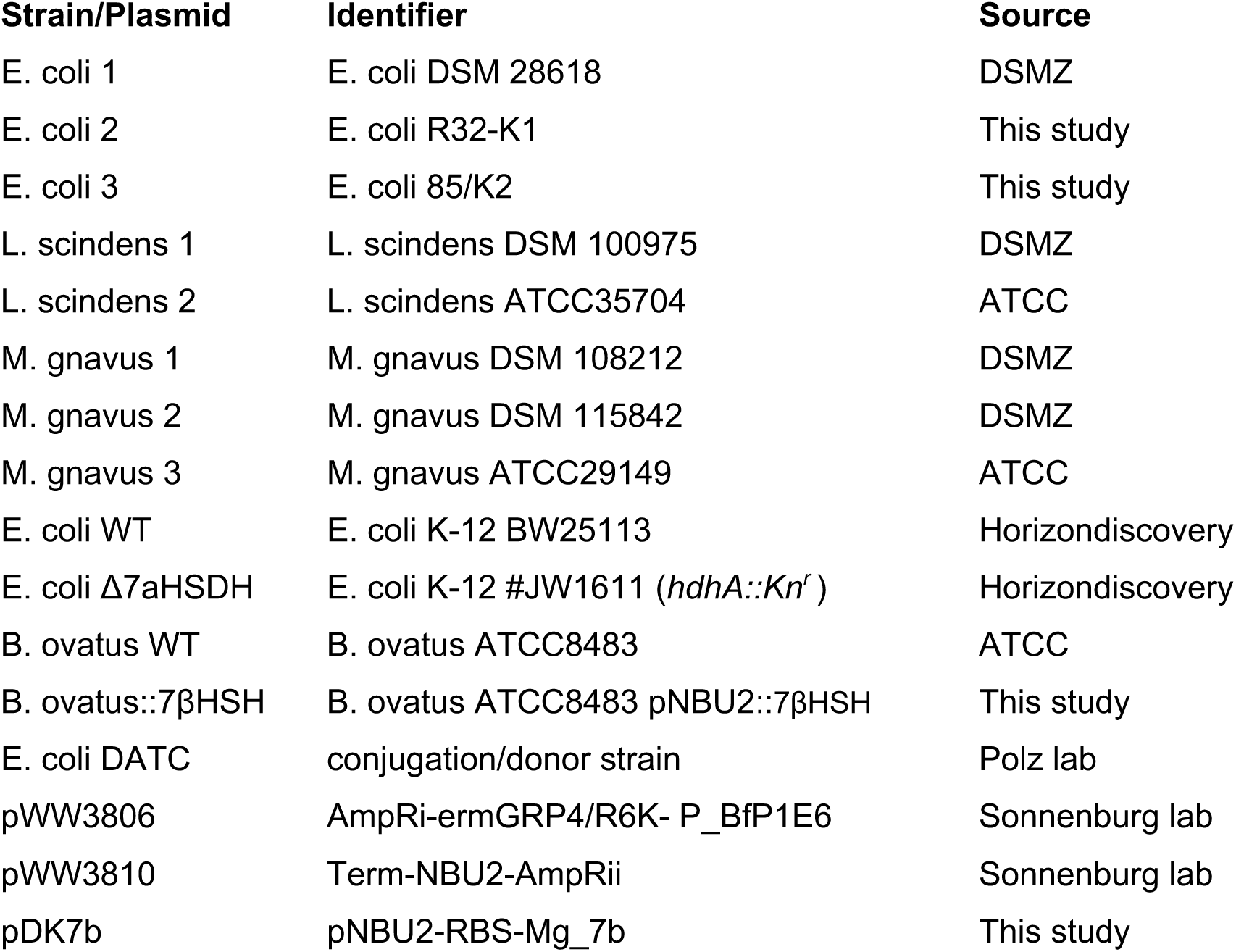

### Organoid culture

Biopsies were collected during endoscopy in DMEM supplemented with HEPES, Pen/Strep, Primocin and RhoK (Y-27632) inhibitor. Epithelial crypts were subsequently isolated to establish patient-derived organoid cultures adapting previously described protocols^66^. In short, the biopsies were cut into small pieces, and incubated in chelating buffer (5.6 mM Na2HPO4, 8 mM Kh2PO4, 96 mM NaCl, 1.6 mM KCl, 44 mM sucrose and 54.8 mM D-sorbitol) supplemented with 6.25 mM DTT and 10 mM EDTA for 30 min at 4°C on a rotating wheel. Crypt structures were squeezed out of the tissue pieces by placing glass slide on top of them and applying gently pressure. These structures were then embedded in basement matrix extract (BME) and cultured in previously described colon medium. Growing 3D organoids were then sheared every week for passaging, freezing or establishing polarized monolayer cultures. For the monolayer cultures, confluent organoids grown on a 12-well plate were collected, sheared using a narrowed Pasteur pipette and two thirds were then seeded on cellQUART 24-Well Cell Culture Inserts (PET clear, 0.4µm, SABEU, Northeim, Germany) which wer pre-coated with for 1 min with a 20% BME solution. Confluent monolayers were observed after three days when the medium was exchanged to differentiation conditions^67^ (hENR: EGF, Noggin, R-sponding media for 5 days) before stimulation experiments were performed.

Large and small intestinal organoid monolayers were treated for 24 hours, basolaterally and apically with 150 μM CDCA, 10 μM GW4064 or DMSO. RNA was extracted using Qiagen RNeasy mini kit, followed by commercial RNA sequencing at Novogene (poly A enrichment, NovaSeq X Plus PE150, 9G raw data per sample). Data was processed using nf-core/rnaseq version 3.14^68^. FGF19 from the basolateral side of small intestinal organoid monolayers was quantified using ELISA (Quantikine).

### scRNA analysis

On a cohort of UC patients used for fecal bile acid analysis, we carried out colonic biopsy single-cell RNA-seq (38 biopsies spanning different endoscopic Mayo scores from 10 patients, subset of Kohl et al, currently in revision). Briefly, mucosal biopsies were obtained during routine colonoscopy from multiple regions of the distal colon and rectum. Biopsies were immediately placed in supplemented DMEM containing antibiotics and ROCK inhibitor and processed fresh. After washing, epithelial cells were separated by EDTA incubation and mechanical dissociation, while the remaining tissue fraction was enzymatically digested with Liberase and DNase I. Epithelial and non-epithelial fractions were then combined, subjected to red blood cell lysis, Fc blocking, and stained with antibodies against EPCAM, CD45, viability dye, and sample-specific TotalSeq-C hashtag antibodies to enable multiplexing of biopsy regions. Live cells were flow-sorted into epithelial, immune, and stromal-enriched fractions based on EPCAM and CD45 expression, pooled by patient, and processed using the 10x Genomics Chromium Single Cell 5′ platform with matched gene-expression and hashtag libraries. Sequencing reads were aligned to the human genome using Cell Ranger, ambient RNA was removed with SoupX, and downstream analysis was performed in Seurat (version 4.1.1). Cells were filtered for quality based on detected gene number (200-2500 mapped features), mitochondrial read fraction (<25% mapped reads), and unambiguous hashtag assignment to retain singlets. Data were log-normalized (LoNormalize method with scale factor 10,000), clustered (dimension=50), and visualized by UMAP. Broad epithelial, immune, and stromal populations were first identified using canonical marker genes, followed by separate reclustering of each compartment and annotation of refined cell types based on established cell-type-specific expression patterns. Differential marker genes were identified using Seurat’s FindAllMarkers function and used to support cell type assignments.

Individual BA clusters were centered log-ratio (CLR) transformed, and cell compositions at each pre-specified hierarchy node were represented using isometric log-ratio (ILR) coordinates as described above. Branch-level R^2^ values were calculated as variance-weighted averages across cell composition ILR coordinates. To identify BA-associated mucosal cell state remodeling, we performed a pre-specified hierarchical composition screen focused on host-derived BA Cluster a and epimerized host-derived BA Cluster b. At each internal node of the cell-state hierarchy, ILR-transformed child compositions were modeled as a function of BA cluster CLR and Mayo score. Incremental BA-associated variance was calculated as: ΔR_BA_^2^=R^2^_(Mayo+BA cluster)_-R^2^_(Mayo)_. Coordinate-level P values were combined across ILR axes using Fisher’s method and Benjamini–Hochberg corrected across tested hierarchy branches. Secretory epithelial subtype associations were further summarized using patient-level Spearman correlations between within-secretory subtype fractions and BA cluster CLR values. Selected biopsy-level cell-state associations were analyzed using within-parent cell fractions. Biopsy-level sensitivity models used random-effects regression with patient as a random intercept and Mayo score as covariate. Goblet-cell maturation-state enrichment was tested at the patient level using quasibinomial models of each state count relative to the total goblet maturation arc count as a function of fecal Cluster a CLR. A biopsy-level Mayo-adjusted random-effects sensitivity analysis was performed for each maturation state using patient as a random intercept. For transcriptomic pathway analysis across the goblet maturation arc, patient-level pseudobulk profiles were generated by summing raw counts across cells from the specified maturation state or the whole maturation arc. Counts were normalized with edgeR and modeled with limma-voom. Genes were ranked by the moderated t-statistic for association with fecal Cluster a CLR. Reactome pathway enrichment was tested using fgsea. Histological validation was performed in an independent UC cohort with matched stool BA profiles and PAS-stained colonic biopsies (n=14)^28^. PAS-stained sections were used to quantify goblet-cell density normalized to crypt length (average 5 field of view / biopsy scan with proper crypt orientation). Patients were stratified by fecal host-derived BA Cluster a abundance above or below 10%. Group differences were assessed using the Mann–Whitney U test, and associations with fecal Cluster a abundance were assessed using Spearman correlation.

### TR-FRET

BA were tested for their ability to activate FXR in a cell-free fluorescence resonance electron transfer (FRET) assay using LanthaScreen™ technology according to the manufacturer’s protocol (Thermo Fisher™, PV4833). Fluorescence detection was done in a Tecan SPARK Microplate reader (Tecan Group, Switzerland) set up according to the LanthaScreen™ Terbium Assay Setup guide available at www.lifetechnologies.com/instrumentsetup. Results were expressed as ratio of fluorescence at 520 nm/485 nm.

### Quantification of bile acids and 7α-hydroxy-4-cholesten-3-one

BA analysis of validation cohort stool samples and bacterial culture supernatants was performed as previously described^30^. Briefly, stool samples were dried in a vacuum centrifuge. Ten milligrams of dried stool was weighted into a Lysing Matrix E tube and subjected to bead-beating at 6000 rpm for 3 × 30 seconds. After homogenization, 0.2 mL of 50 mM ammonium acetate (pH 5.6) and 0.5 mL acetonitrile were added to the samples, and they were put for 1 hour in a shaking heating block at 60°C. Samples were centrifuged and samples were subjected to a second round of extraction with 0.3 mL methanol and 0.5 mL isopropanol. Combined supernatants were taken for quantification of BAs. BAs and their conjugates were quantified using liquid chromatography–tandem mass spectrometry, employing selected reaction monitoring. In brief, 1 μL of the extract was injected on a Kinetex C8 column (100 Å, 100 × 2.1 mm; Phenomenex) with the respective guard column, employing a flow rate of 130 μL/min. After loading at 80% A (2.5 mM ammonium acetate in water) a 20-minute-long gradient from 60% B to 90% B (2.5 mM ammonium acetate in methanol) was used for separation. The high-performance liquid chromatography (RSLC ultimate 3000; Thermo Fisher Scientific) was directly coupled to a TSQ Quantiva mass spectrometer (Thermo Fisher Scientific) via electrospray ionization. BAs and their conjugates were analyzed in the negative ion mode, employing the respective transitions (e.g. glycocholic acid *m/z* 464 to *m/z* 73) and an optimized collision energy, which has been determined by analyzing authentic standards. Chromatograms have been manually interpreted using trace finder (Thermo Fisher Scientific), validating experimental retention times with the respective quality controls of the pure substances. 7α-hydroxy-4-cholesten-3-one (C4) was measured from patient serum samples using identical chromatographic conditions with the transitions *m/z* 401 to 144 *<u>m/z</u>* and *m/z* 401 to *m/z* 177 in the positive ion mode.

### Gut-liver feedback axis analysis

Publicly available iHMP ileal biopsy host RNA-seq count data, biopsy 16S profiles, and matched stool bile acid metadata were analyzed for samples with available transcriptomic and metabolomic measurements (https://ibdmdb.org/). Crohn’s disease samples were excluded for analyses to reduce confounding from ileal inflammation, non-IBD controls were analyzed separately where indicated. Samples were stratified by fecal host-derived bile acid abundance using the predefined high/low HBA threshold of 10%. Host RNA-seq count matrices were filtered to retain samples with at least 1,000 total counts and 500 detected genes, and genes with counts ≥10 in at least three samples. FXR activity in iHMP ileal biopsies was inferred using single-sample GSEA from the GSVA package. A composite FXR activity score was calculated as the z-scored FXR-up score minus the z-scored FXR-down score. Differential expression between high- and low-HBA groups was performed on variance-stabilized expression values using limma. Genes were ranked by moderated t-statistic, and enrichment of FXR-up and FXR-down signatures was tested using fgsea with 10,000 permutations. FGF19 gene expression was extracted from the normalized ileal RNA-seq matrix. For microbiome analyses, genus-level 16S profiles were quality controlled by total read count, number of detected taxa, dominance of the most abundant genus, and Shannon diversity. Genera present in at least 10% of samples were retained and transformed using centered log-ratio transformation after adding a pseudocount. Bacterial signatures derived from our taxonomic IBD/BAD meta-analysis (Fig2f) were calculated as the mean CLR abundance of genera assigned to that cluster. Associations between bile acids, inferred FXR activity, FGF19 expression, C4 and bacterial signatures were tested using Spearman correlation.

## Data and code availability

Data tables and scripts necessary to generate figures and tables from this manuscript have been deposit at figshare. Tables including patient metadata are available upon reasonable request. Raw organoid transcriptomics and MBRA 16S-rRNA amplicon sequencing data has been deposited at GEO and ENA. Bioinformatical pipelines to predict BA composition from metagenomics and quantify our described BA converting enzyme modules as well as BA cluster are publicly available at github.com/MaximilianBaumgartner/MetaBile.

## Ethical approval

Approval to collect patient material in our confirmation cohorts was granted by the responsible ethics commissions, for BA analysis in the scRNA cohort and organoid isolation (Medical University of Vienna EK 1260/2022), BA analysis in the FGF19 and C4 confirmation cohort (Medical University of Graz EK 34-463 ex 21/22 1263-2022), histological confirmation and bacterial strain isolation cohort (Medical University of Vienna EK 1617/2014, 1780/2019, 1910/2019).

## References

1. Kaplan, G. G. & Windsor, J. W. The four epidemiological stages in the global evolution of inflammatory bowel disease. Nat. Rev. Gastroenterol. Hepatol. 18, 56–66 (2021).

2. Raine, T. & Danese, S. Breaking through the therapeutic ceiling: What will it take? Gastroenterology 162, 1507–1511 (2022).

3. Alsoud, D., Verstockt, B., Fiocchi, C. & Vermeire, S. Breaking the therapeutic ceiling in drug development in ulcerative colitis. Lancet Gastroenterol. Hepatol. 6, 589–595 (2021).

4. Franzosa, E. A. et al. Gut microbiome structure and metabolic activity in inflammatory bowel disease. Nature Microbiology 4, 293–305 (2019).

5. Imhann, F. et al. Interplay of host genetics and gut microbiota underlying the onset and clinical presentation of inflammatory bowel disease. Gut 67, 108–119 (2018).

6. Zheng, J. et al. Noninvasive, microbiome-based diagnosis of inflammatory bowel disease. Nat. Med. 30, 3555–3567 (2024).

7. Iliev, I. D., Ananthakrishnan, A. N. & Guo, C.-J. Microbiota in inflammatory bowel disease: mechanisms of disease and therapeutic opportunities. Nat. Rev. Microbiol. 23, 509–524 (2025).

8. Fishbein, S. R. S., Mahmud, B. & Dantas, G. Antibiotic perturbations to the gut microbiome. Nat. Rev. Microbiol. 21, 772–788 (2023).

9. Vich Vila, A., et al. Faecal metabolome and its determinants in inflammatory bowel disease. Gut gutjnl-2022-328048 (2023).

10. Wahlström, A., Sayin, S. I., Marschall, H. U. & Bäckhed, F. Intestinal Crosstalk between Bile Acids and Microbiota and Its Impact on Host Metabolism. Cell Metab. 24, 41–50 (2016).

11. Fuchs, C. D. et al. Bile acid metabolism and signalling in liver disease. J. Hepatol. 82, 134–153 (2025).

12. Hegyi, P., Maléth, J., Walters, J. R., Hofmann, A. F. & Keely, S. J. Guts and Gall: Bile Acids in Regulation of Intestinal Epithelial Function in Health and Disease. Physiol. Rev. 98, 1983–2023 (2018).

13. Hofmann, A. F. Enterohepatic circulation of bile acids. in Handbook of physiology (ed. Bb, S. S. F. J.) vol. 3 567–588 (Oxford University Press, New York, 1989).

14. Dawson, P. A. & Karpen, S. J. Intestinal transport and metabolism of bile acids. J. Lipid Res. 56, 1085–1099 (2015).

15. Ridlon, J. M. & Gaskins, H. R. Another renaissance for bile acid gastrointestinal microbiology. Nat. Rev. Gastroenterol. Hepatol. 21, 348–364 (2024).

16. Mohanty, I. et al. The underappreciated diversity of bile acid modifications. Cell 187, 1801–1818.e20 (2024).

17. Quinn, R. A. et al. Global chemical effects of the microbiome include new bile-acid conjugations. Nature 579, 123–129 (2020).

18. Campbell, C. et al. Bacterial metabolism of bile acids promotes generation of peripheral regulatory T cells. Nature 581, 475–479 (2020).

19. Tanaka, H. et al. Ligand-independent activation of the glucocorticoid receptor by ursodeoxycholic acid. Repression of IFN-gamma-induced MHC class II gene expression via a glucocorticoid receptor-dependent pathway. J. Immunol. 156, 1601–1608 (1996).

20. Peterson, D., Weidenmaier, C., Timberlake, S. & Gura Sadovsky, R. Depletion of key gut bacteria predicts disrupted bile acid metabolism in inflammatory bowel disease. Microbiol. Spectr. 13, e0199924 (2025).

21. Lloyd-Price, J. et al. Multi-omics of the gut microbial ecosystem in inflammatory bowel diseases. Nature (2019) doi:10.1038/s41586-019-1237-9.

22. Lavelle, A. & Sokol, H. Gut microbiota-derived metabolites as key actors in inflammatory bowel disease. Nat. Rev. Gastroenterol. Hepatol. 17, 223–237 (2020).

23. Schirmer, M. et al. Compositional and Temporal Changes in the Gut Microbiome of Pediatric Ulcerative Colitis Patients Are Linked to Disease Course. Cell Host Microbe 24, 600–610.e4 (2018).

24. Jia, B., Park, D., Hahn, Y. & Jeon, C. O. Metagenomic analysis of the human microbiome reveals the association between the abundance of gut bile salt hydrolases and host health. Gut Microbes 11, 1300–1313 (2020).

25. Jin, W.-B. et al. Microbiota-derived bile acids antagonize the host androgen receptor and drive anti-tumor immunity. Cell 188, 2336–2353.e38 (2025).

26. Vital, M., Rud, T., Rath, S., Pieper, D. H. & Schlüter, D. Diversity of Bacteria Exhibiting Bile Acid-inducible 7α-dehydroxylation Genes in the Human Gut. Comput. Struct. Biotechnol. J. 17, 1016–1019 (2019).

27. Li, W. et al. A bacterial bile acid metabolite modulates Treg activity through the nuclear hormone receptor NR4A1. Cell Host Microbe 29, 1366–1377.e9 (2021).

28. Yao, L. et al. A selective gut bacterial bile salt hydrolase alters host metabolism. Elife 7, (2018).

29. Hall, A. B. et al. A novel Ruminococcus gnavus clade enriched in inflammatory bowel disease patients. Genome Med. (2017) doi:10.1186/s13073-017-0490-5.

30. Baumgartner, M. et al. Mucosal Biofilms Are an Endoscopic Feature of Irritable Bowel Syndrome and Ulcerative Colitis. Gastroenterology 161, 1245–1256.e20 (2021).

31. Vijayvargiya, P. et al. Analysis of fecal primary bile acids detects increased stool weight and colonic transit in patients with chronic functional diarrhea. Clin. Gastroenterol. Hepatol. 17, 922–929.e2 (2019).

32. Camilleri, M. et al. Comparison of biochemical, microbial and mucosal mRNA expression in bile acid diarrhoea and irritable bowel syndrome with diarrhoea. Gut gutjnl-2022-327471 (2022).

33. Hillman, E. B. M. et al. Ruminococcus gnavus and biofilm markers in feces from primary bile acid diarrhea patients indicate new disease mechanisms and potential for diagnostic testing. Gastro Hep Adv. 4, 100712 (2025).

34. Auchtung, J. M., Robinson, C. D., Farrell, K. & Britton, R. A. MiniBioReactor arrays (MBRAs) as a tool for studying C. difficile physiology in the presence of a complex community. Methods Mol. Biol. 1476, 235–258 (2016).

35. Camilleri, M., Carlson, P., Acosta, A. & Busciglio, I. Colonic mucosal gene expression and genotype in irritable bowel syndrome patients with normal or elevated fecal bile acid excretion. Am. J. Physiol. Gastrointest. Liver Physiol. 309, G10–20 (2015).

36. Damianos, J., Abdelnaem, N. & Camilleri, M. Gut goo: Physiology, diet, and therapy of intestinal mucus and biofilms in gastrointestinal health and disease. Clin. Gastroenterol. Hepatol. 23, 205–215 (2025).

37. Devlin, A. S. & Fischbach, M. A. A biosynthetic pathway for a prominent class of microbiota-derived bile acids. Nat. Chem. Biol. (2015) doi:10.1038/nchembio.1864.

38. Best, W. R., Becktel, J. M., Singleton, J. W. & Kern, F., Jr. Development of a Crohn’s disease activity index. National Cooperative Crohn’s Disease Study. Gastroenterology 70, 439–444 (1976).

39. Schroeder, K. W., Tremaine, W. J. & Ilstrup, D. M. Coated oral 5-aminosalicylic acid therapy for mildly to moderately active ulcerative colitis. A randomized study. N. Engl. J. Med. 317, 1625–1629 (1987).

40. van der Post, S. et al. Structural weakening of the colonic mucus barrier is an early event in ulcerative colitis pathogenesis. Gut 68, 2142–2151 (2019).

41. Singh, V. et al. Chronic inflammation in ulcerative colitis causes long-term changes in goblet cell function. Cell. Mol. Gastroenterol. Hepatol. 13, 219–232 (2022).

42. Vítek, L. Bile acid malabsorption in inflammatory bowel disease. Inflamm. Bowel Dis. 21, 476–483 (2015).

43. Mueller, M. et al. Ursodeoxycholic acid exerts farnesoid X receptor-antagonistic effects on bile acid and lipid metabolism in morbid obesity. J. Hepatol. 62, 1398–1404 (2015).

44. Gothe, F. et al. Bile acid malabsorption assessed by 7 alpha-hydroxy-4-cholesten-3-one in pediatric inflammatory bowel disease: correlation to clinical and laboratory findings. J. Crohns. Colitis 8, 1072–1078 (2014).

45. Battat, R. et al. Serum Concentrations of 7α-hydroxy-4-cholesten-3-one Are Associated With Bile Acid Diarrhea in Patients With Crohn’s Disease. Clin. Gastroenterol. Hepatol. 17, 2722–2730.e4 (2019).

46. Reiman, D., Layden, B. T. & Dai, Y. MiMeNet: Exploring microbiome-metabolome relationships using neural networks. PLoS Comput. Biol. 17, e1009021 (2021).

47. Topçuoðlu, B. D., Lesniak, N. A., Ruffin, M. T., 4th, Wiens, J. & Schloss, P. D. A framework for effective application of machine learning to microbiome-based classification problems. MBio 11, e00434–20 (2020).

48. Baumgart, M. et al. Culture independent analysis of ileal mucosa reveals a selective increase in invasive Escherichia coli of novel phylogeny relative to depletion of Clostridiales in Crohn’s disease involving the ileum. ISME J. 1, 403–418 (2007).

49. Smith, P. & Bénézech, C. Creeping fat in Crohn’s disease: Innocuous or innocuum? Immunity vol. 53 905–907 (2020).

50. Holani, R. et al. Bile acid-induced metabolic changes in the colon promote Enterobacteriaceae expansion and associate with dysbiosis in Crohn’s disease. Sci. Signal. 17, eadl1786 (2024).

51. Sinha, S. R. et al. Dysbiosis-Induced Secondary Bile Acid Deficiency Promotes Intestinal Inflammation. Cell Host Microbe 27, 659–670.e5 (2020).

52. Sayin, S. I. et al. Gut microbiota regulates bile acid metabolism by reducing the levels of tauro-beta-muricholic acid, a naturally occurring FXR antagonist. Cell Metab. 17, 225–235 (2013).

53. Hofmann, A. F. The enterohepatic circulation of bile acids in mammals: form and functions. Front. Biosci. (Landmark Ed.) 14, 2584–2598 (2009).

54. Kim, C. Y. et al. Human reference gut microbiome catalog including newly assembled genomes from under-represented Asian metagenomes. Genome Med. 13, 134 (2021).

55. Kaminski, J. et al. High-specificity targeted functional profiling in microbial communities with ShortBRED. PLoS Comput. Biol. 11, e1004557 (2015).

56. Abramson, J. et al. Accurate structure prediction of biomolecular interactions with AlphaFold 3. Nature 630, 493–500 (2024).

57. van Kempen, M. et al. Fast and accurate protein structure search with Foldseek. Nat. Biotechnol. 42, 243–246 (2024).

58. Blanco-Míguez, A. et al. Extending and improving metagenomic taxonomic profiling with uncharacterized species using MetaPhlAn 4. Nat. Biotechnol. 41, 1633–1644 (2023).

59. Beghini, F. et al. Integrating taxonomic, functional, and strain-level profiling of diverse microbial communities with bioBakery 3. Elife 10, (2021).

60. Buchfink, B., Xie, C. & Huson, D. H. Fast and sensitive protein alignment using DIAMOND. Nat. Methods 12, 59–60 (2015).

61. Mallick, H. et al. Multivariable association discovery in population-scale meta-omics studies. PLoS Comput. Biol. 17, e1009442 (2021).

62. Rytter, H. et al. In vitro microbiota model recapitulates and predicts individualised sensitivity to dietary emulsifier. Gut 74, 761–774 (2025).

63. Callahan, B. J. et al. DADA2: High-resolution sample inference from Illumina amplicon data. Nat. Methods 13, 581–583 (2016).

64. Lundin, D. & Andersson, A. SBDI Sativa curated 16S GTDB database. Linnéuniversitetet 10.17044/SCILIFELAB.14869077 (2026).

65. Bobonis, J., Yang, A. L. J., Voogdt, C. G. P. & Typas, A. TAC-TIC, a high-throughput genetics method to identify triggers or blockers of bacterial toxin-antitoxin systems. Nat. Protoc. 19, 2231–2249 (2024).

66. Sato, T. et al. Long-term expansion of epithelial organoids from human colon, adenoma, adenocarcinoma, and Barrett’s epithelium. Gastroenterology 141, 1762–1772 (2011).

67. Martinez-Silgado, A., Beumer, J. & Clevers, H. Directed differentiation of Murine and human small intestinal organoids toward all mature lineages. Methods Mol. Biol. 2650, 107–122 (2023).

68. Ewels, P. A. et al. The nf-core framework for community-curated bioinformatics pipelines. Nat. Biotechnol. 38, 276–278 (2020).

