## Extended data Figures S1-7 for "Host-derived bile acids drive dysbiosis by selecting bile-resistant epimerizing bacteria in inflammatory bowel disease"

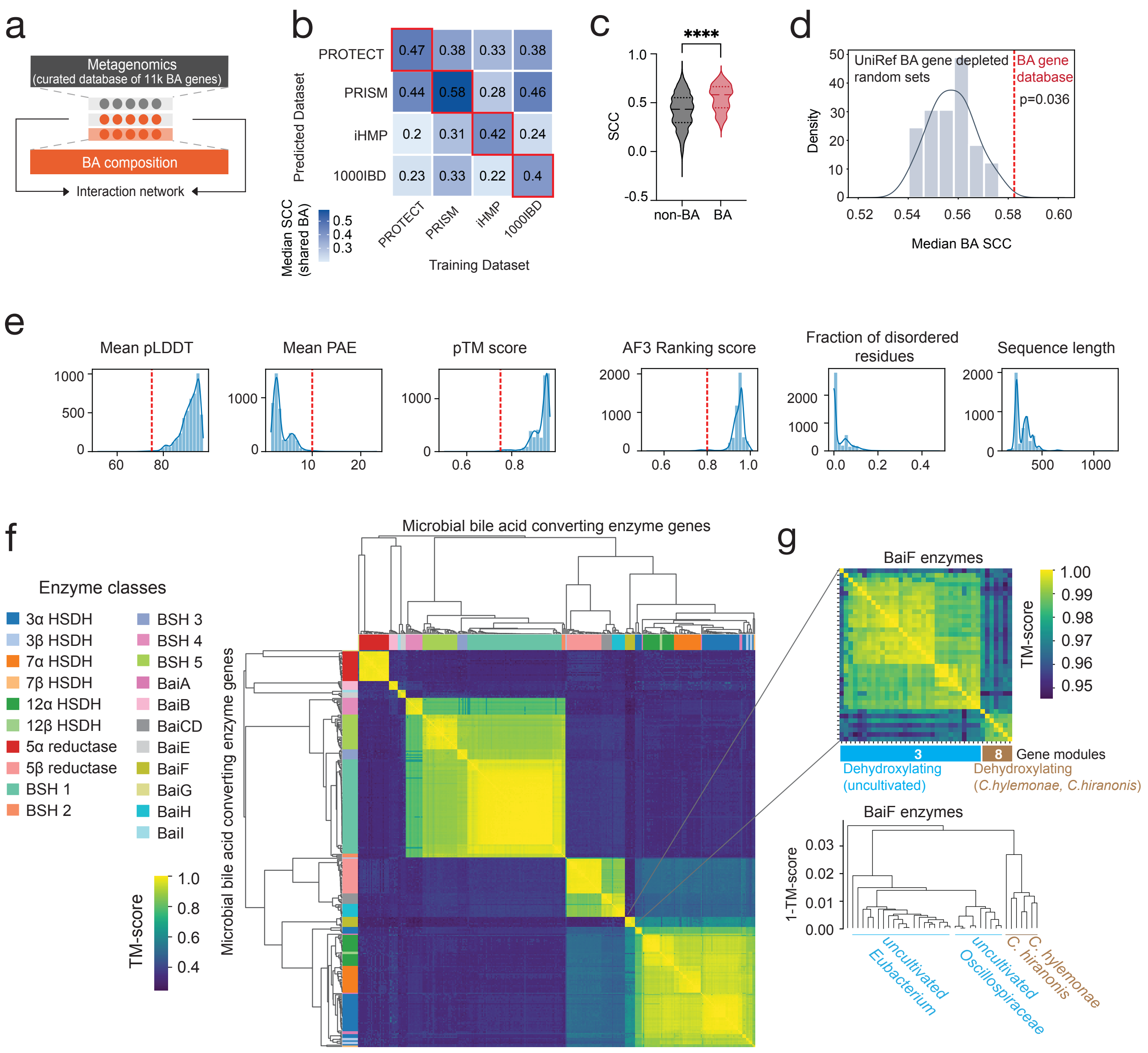


**Extended Data Fig. 1: Validation of neural network approach linking microbial bile acid genes and stool bile acid composition.** (**a**) Stool metagenomic sequencing reads from four inflammatory bowel disease datasets (PROTECT n= 88, PRISM n= 214, iHMP n= 459, 1000IBD n= 355) were mapped to a custom microbial bile acid (BA) converting enzyme gene database (11,465 bile-acid-transforming enzyme sequences spanning bile salt hydrolases (BSHs), hydroxysteroid dehydrogenases (HSDHs), *Bai* operon genes A–I (C7 dehydroxylation) and C5 reductases using the shortBRED package (see **Methods** and github:Maximilian_Baumgartner/metaBile). Neural networks were trained to predict BA composition in paired metagenomics-metabolomics samples using the MiMeNet package (see **Methods**). (**b**) Median Spearman correlation coefficients (SCC) between predicted and observed BAs including intra- (red border) and across-datasets. Across-datasets models were trained only on BAs being present in both datasets. (**c**) SCC of BA and non-BA features in the main cohort used for BA cluster definition (PRISM). Statistical significance determined with a Mann-Whitney U test,****p<0.0001. (**d**) Distribution of median SCC for BA prediction by BA-related and unrelated microbial genes. MiMeNet was trained on 50 prevalence/abundance-matched BA enzyme-depleted random gene sets versus our custom BA converting enzyme gene database using the PRISM dataset. (**e**) Representative BA-modifying proteins were modeled with AlphaFold3. Sequences were clustered at 85% amino-acid identity, yielding 4,866 representative proteins for structural validation. Structures were filtered to retain high-confidence models using the following criteria: mean pLDDT > 75, mean PAE < 10 Å, pTM > 0.75 and ranking_score > 0.8. (**f**) Pairwise structural similarities were computed in an all-against-all comparison using Foldseek in TM-align mode, and hierarchical clustering was performed on the resulting TM-score distance matrix (1 − TM-score) using the Ward method. The number of BSH structural clusters was selected using the maximum silhouette criterion. (**g**) Example of structural heterogeneity among BaiF proteins. Top, TM-score heatmap; bottom, hierarchical clustering dendrogram based on 1 − TM-score. BaiF proteins from uncultured bacteria detected only by metagenomics formed a distinct structural group from BaiF proteins of *Clostridium hylemonae* and *Clostridium hiranonis*.

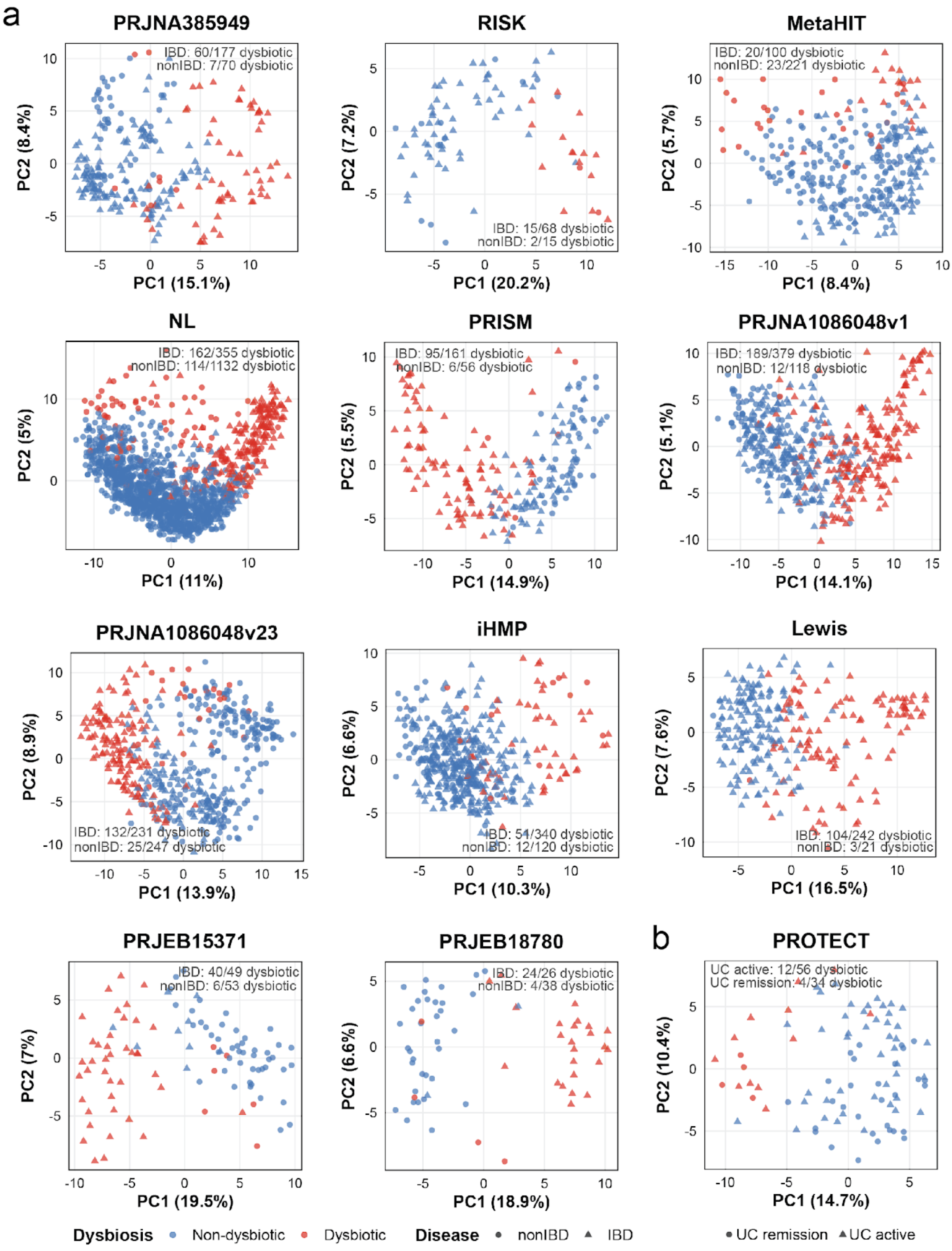


**Extended Data Fig. 2: Definition and validation of dysbiosis scores across meta-analysis cohorts.** To enable microbiome-state comparisons across cohorts, taxonomic profiles from high-quality metagenomic reads were recalculated using a MetaPhlAn4-based pipeline (see **Methods**). Inflammatory bowel disease associated (IBD) dysbiosis scores were computed using median Bray–Curtis dissimilarity to non-IBD reference samples, with dysbiosis defined by the 90th percentile threshold of the non-IBD reference distribution. Showing genus-level Aitchison distance Principal Coordinates Analysis (PcoA). Dysbiotic samples colored in red, non-dysbiotic samples in blue. (**a**) IBD samples shown as triangles, non-IBD as circles. (**b**) In the pediatric ulcerative colitis cohort PROTECT, clinical remission was used as the closest available low-disease reference state.

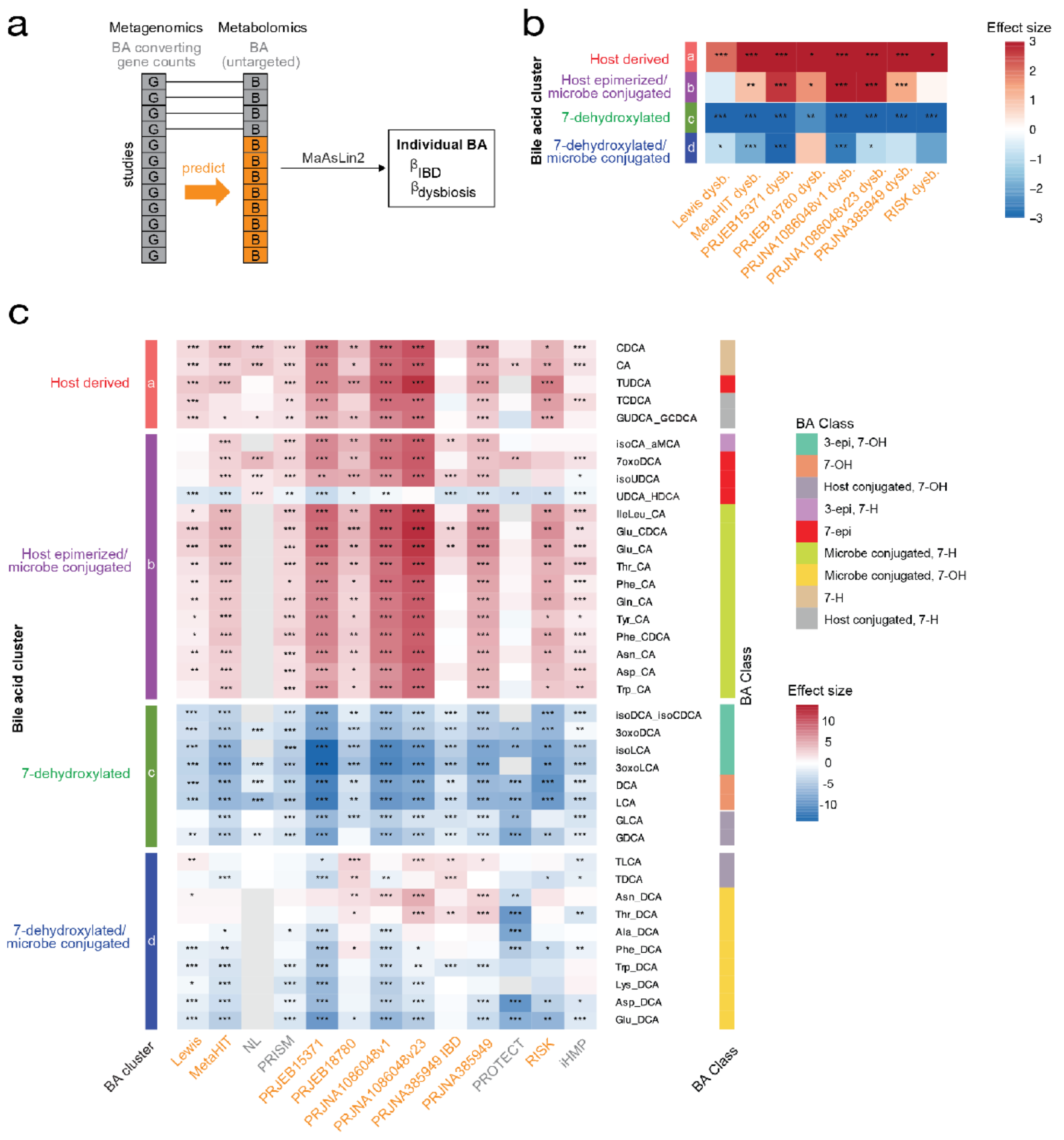


**Extended Data Fig. 3: Meta-analysis of individual bile acid associations across cohorts.**
(**a**) Cohort-specific associations between individual bile acids (BA) and inflammatory bowel disease (IBD)-associated dysbiosis were estimated across 12 datasets using observed BA profiles when available (gray: NL, PRISM, PROTECT, iHMP) or predicted bile acid abundances (orange: Lewis, MetaHIT, PRJEB15371, PRJEB18780, PRJNA1086048v1/23, PRJNA385949 and RISK). (**b**) Analyses of predicted BA clusters in multiple IBD cohorts. (**c**) Heatmap summary of effect sizes per cohort. Rows ordered by neural-network-defined BA cluster and bile acid enzymatic class; columns grouped by dysbiosis vs. eubiosis comparison in indicated study. Tile color indicates the direction and magnitude of the association (log2 fold change), asterisks indicate significance after multiple-testing correction within cohort-level analysis. Statistics: Per-cohort BA associations were estimated using MaAsLin2, with subject included as a random effect in longitudinal datasets. Multiple-testing correction using Benjamini–Hochberg FDR within each cohort-level analysis; asterisks denote significant FDR q value, *<0.05, **<0.01, and ***<0.001. Sample size and epidemiological data of all analyzed cohorts in **Supplementary Table 1**.

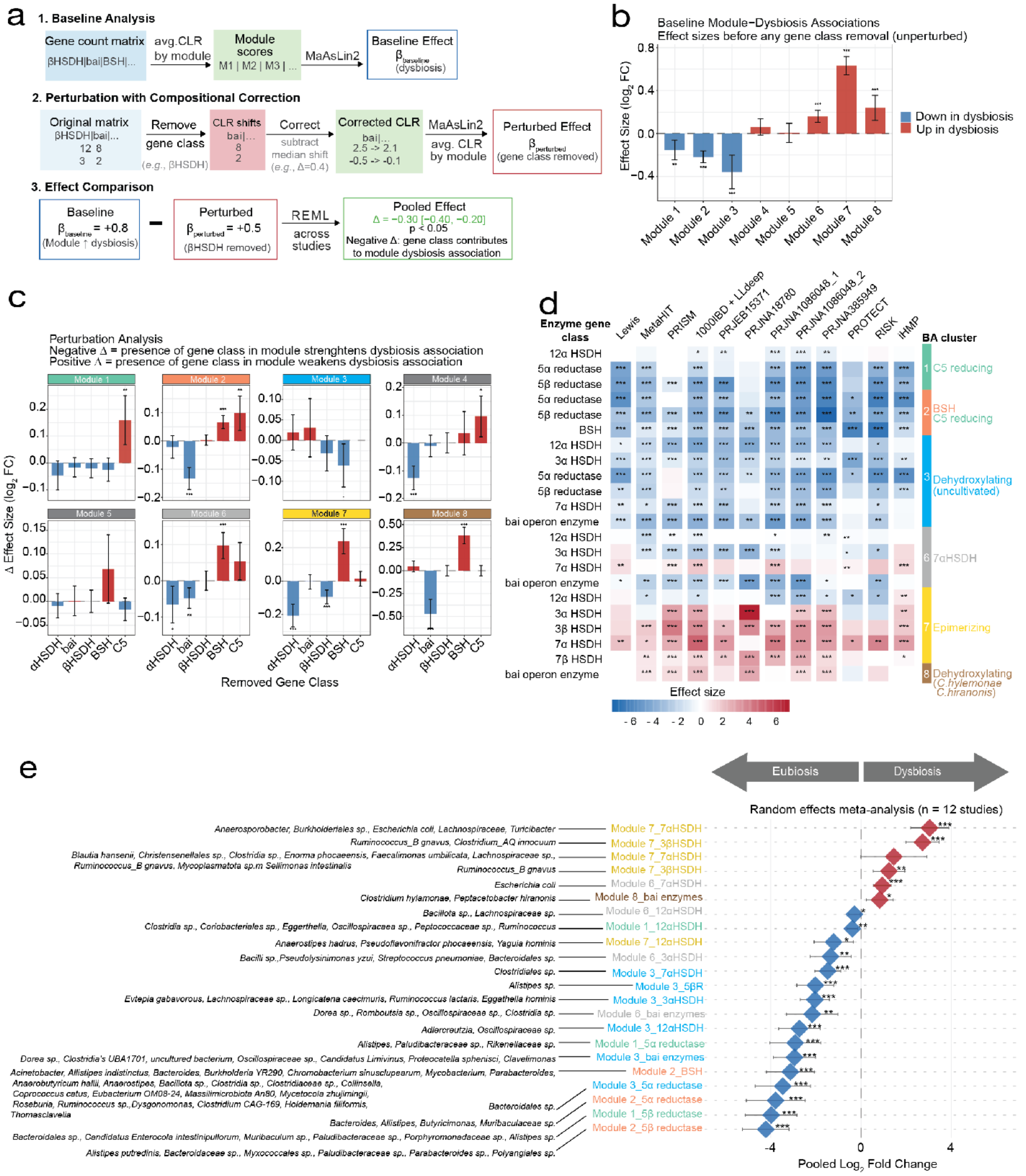


**Extended Data Fig. 4: In silico perturbation analysis identifies bile acid enzyme classes contributing to gene module–dysbiosis associations.** (**a**) Schematic of in silico perturbation approach. (1.) Baseline associations between neural-network-defined BA enzyme modules and dysbiosis were estimated across cohorts. (2.) Selected enzyme classes (BSH, C5 reductases, bai genes, HSDH types) were computationally removed, remaining counts were compositionally corrected, and module abundances were recalculated. (3.) Changes in module–dysbiosis association after perturbation were compared to the unperturbed baseline to quantify the contribution of each enzyme class. (**b**) Baseline module–dysbiosis associations across cohorts. Bars show meta-analyzed effect sizes for each neural-network-defined module in dysbiotic versus eubiotic samples. (**c**) Perturbation effects on module–dysbiosis associations. Bars show change in effect size (Δ log2 fold change relative to the unperturbed model) after removal of each enzyme class. (**d**,**e**) Validation of the association between selected module–enzyme class combinations and dysbiosis across 12 IBD metagenomics datasets. (**d**) Cohort-specific effects shown as a heatmap. (**e**) Forest plot with pooled random-effects summary and taxonomic classification of individual proteins. Statistics: Study-level module associations were estimated using MaAsLin2, with dysbiosis status as the fixed effect and subject included as a random effect for longitudinal datasets. 'Dysbiotic' was defined as >90$^{th}$ median percentile taxonomic Bray-Curtis dissimilarity to controls. Cross-cohort summary estimates were obtained by random-effects meta-analysis. Effect sizes are shown on a log2 scale; error bars indicate 95% confidence intervals. Multiple-testing correction used the Benjamini–Hochberg false discovery rate (FDR) across tested module–enzyme class combinations, asterisks denote significant FDR q value, *<0.05, **<0.01, and ***<0.001. Sample size and epidemiological data of all analyzed cohorts in **Supplementary Table 1**.

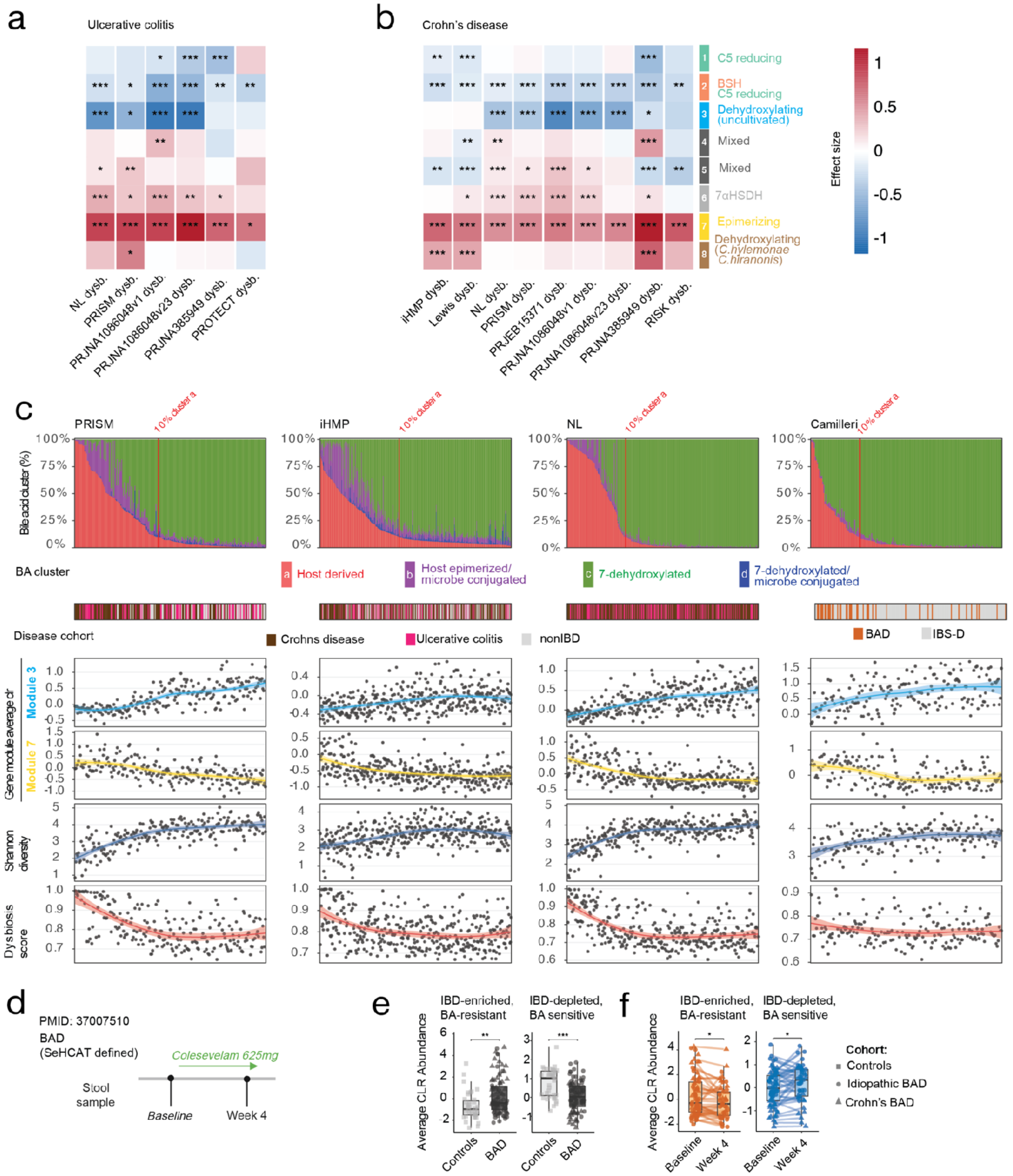


**Extended Data Fig. 5: Subtype-specific meta-analysis of bile acid clusters in IBD, waterfall plots of microbiome variables in main cohorts and effect of colesevelam on bile acid diarrhea-associated dysbiosis.** (**a**,**b**) Validation of the association between IBD-associated dysbiosis and neural-network defined BA converting gene modules in 12 IBD metagenomics datasets, analysis split for IBD subtype (ulcerative colitis and Crohn's disease), columns grouped by study. Tile color indicates the direction and magnitude of the association (log2 fold change), asterisks indicate significance after multiple-testing correction within cohort-level analyses. (**c**) Multi-track waterfall plots of BA and microbiome features in three IBD and one bile acid diarrhea (BAD) cohort. Showing neural-network defined BA cluster abundance (10% relative abundance of host-derived BA cluster a used as a clinical cut-off for bile acid malabsorption, denoted by red line), disease cohort (Crohn's disease, ulcerative colitis, non-IBD controls, IBS-D with or without BAD), BA dehydroxylating gene module 3, BA epimerizing gene module 7, Shannon diversity index and Dysbiosis score. **(d)** Study design of the idiopathic bile acid diarrhoea (BAD) Colesevelam cohort (PMID: 37007510). Stool samples from BAD patients (idiopathic BAD; squares), Crohn's disease with BAD (triangles) and controls (circles) were compared at baseline, and patients with BAD were followed longitudinally after Colesevelam treatment. (**e**) Baseline average centered log-ratio (CLR) abundance of IBD-enriched bile acid-resistant genera (left) and IBD-depleted bile acid-sensitive genera (right) in controls and BAD patients. (**f**) Relative abundance of IBD-enriched bile acid-resistant genera (left) and IBD-depleted bile acid-sensitive genera (right) in paired BAD patient stool samples at baseline and 4 weeks after continuous Colesevelam treatment. Significance in (**a,b**) was assessed using MaAsLin2 with subject included as a random effect in longitudinal datasets and a minimum group size for individual comparisons of n=12, in (**e**) with a Wilcoxon test, and in (**f**) with linear mixed-effects models. Multiple-testing correction using Benjamini–Hochberg FDR within each cohort-level analysis; asterisks denote significant FDR q value, *<0.05, **<0.01, and ***<0.001.

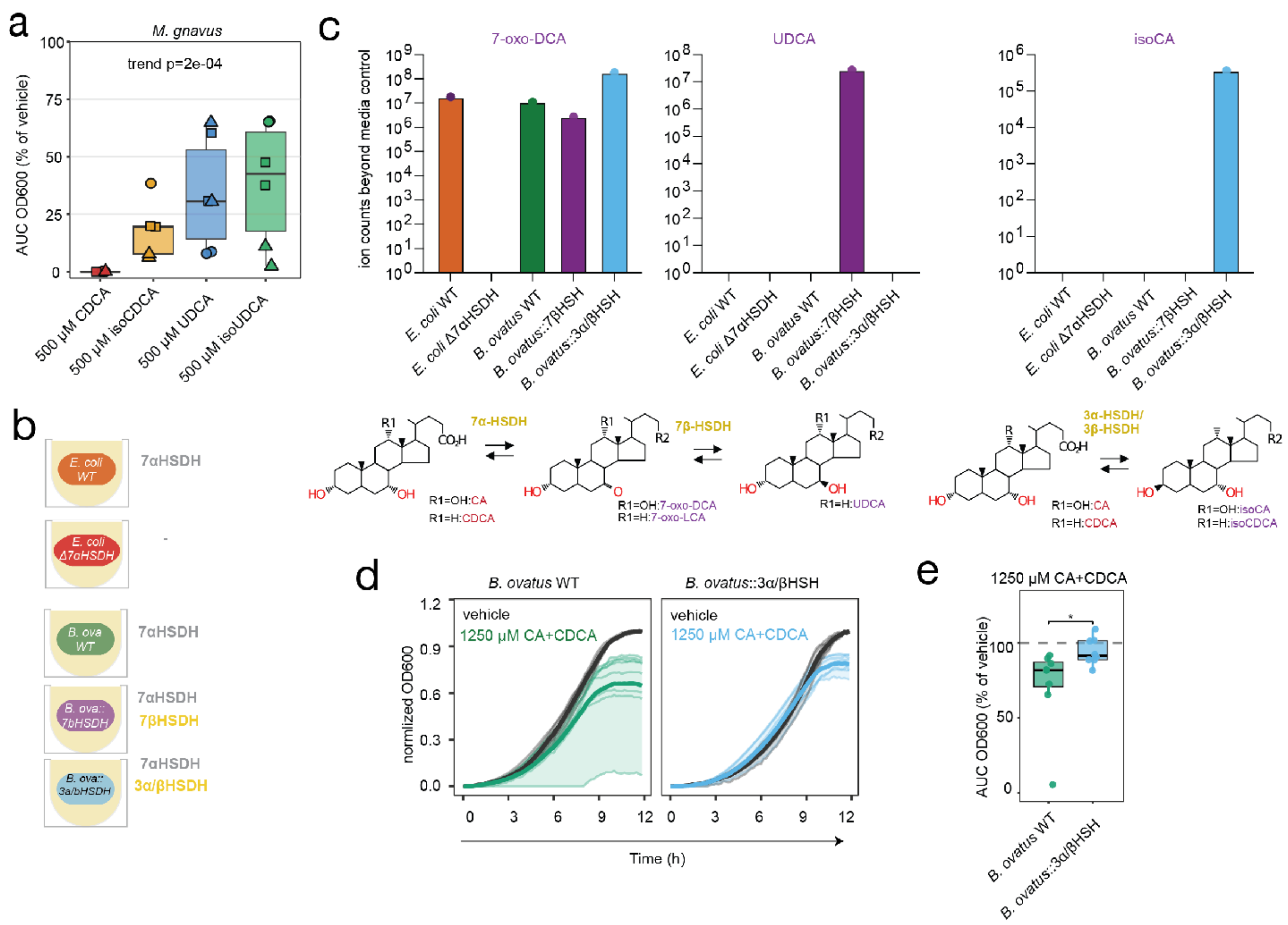


**Extended Data Fig. 6: BA conversion by bacterial strains.** (**a**) *M. gnavus* growth at 500 µM of CDCA, isoCDCA, UDCA or isoUDCA. Showing area under the curve (AUC) for growth measured by OD600 over 12 hours relative to vehicle. **(b)** Depiction of hydroxysteroid-dehydrogenases (HSDHs) present in *E. coli* and *B. ovatus* strains shown in c,d. *E. coli* Δ7αHSDH (*hdhA:: Kn$^r$*) from the Keio collection (orange) and matching WT parental strain (red); *B. ovatus* with genomic integration of 7βHSDH gene (*B. ovatus*::7βHSDH, purple) or 3αHSDH + 3βHSDH (*B. ovatus*::3α/βHSDH blue) from *M. gnavus* and WT parental strain (green). For further details on bacterial strains, see **Methods**. **(c)** HSDH activity in engineered strains. 100 µM of BA precursors (CA/CDCA) were incubated overnight followed by targeted mass spectrometry. (**d,e**) Growth of *B. ovatus*::3a/βHSDH vs. parental WT strain under 1250 µM of CA+CDCA (2:1). Showing growth curves (**d**) and growth relative to vehicle (**e**). Data shown in (**a,d,e**) are representative from two independent experiments. (**a**) 3 independent *M. gnavus* strains in biological replicates. (**d,e**) n=8 biological replicates. Significance was determined with a Jonckheere trend test (**a**) or two-sided Mann-Whitney U test (**e**). *p<0.05.

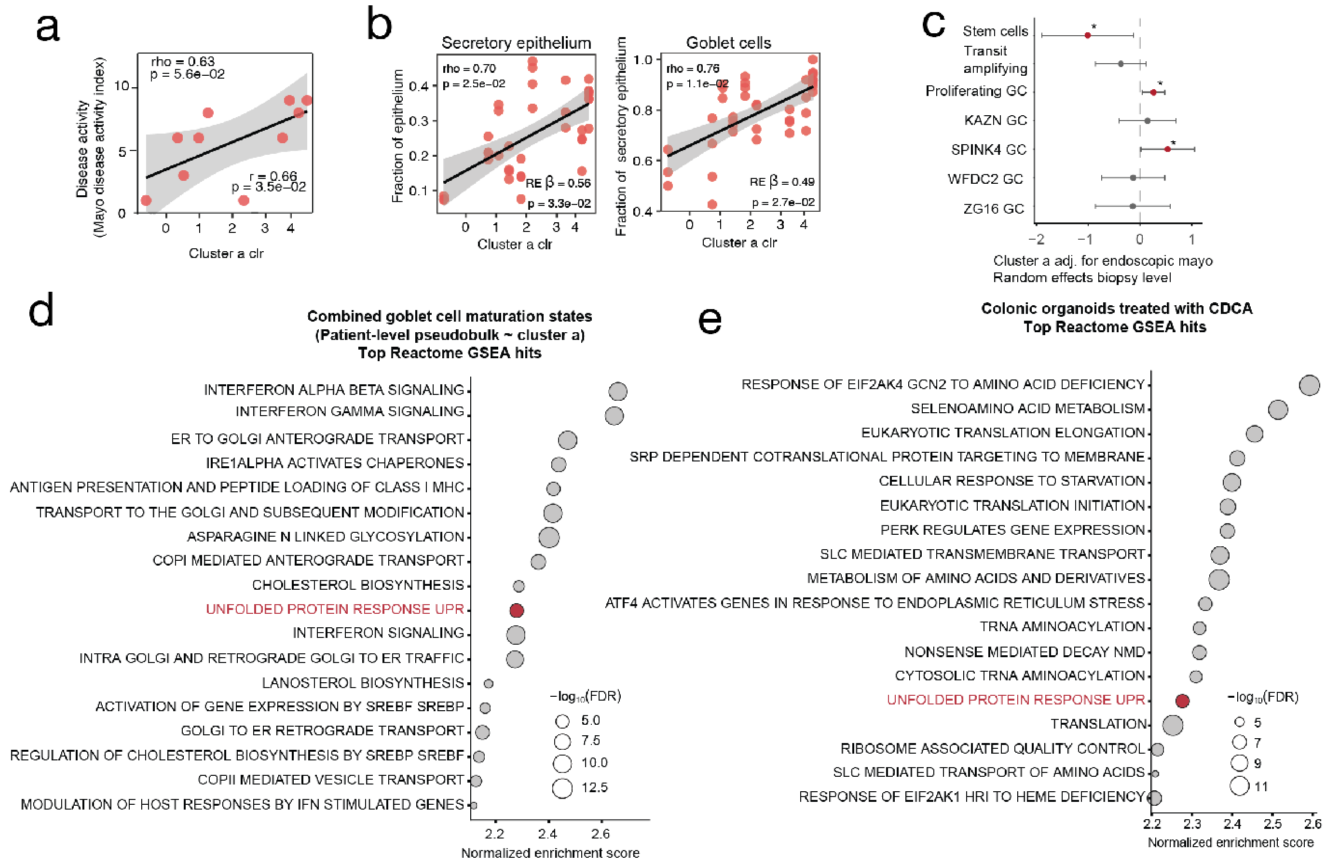


**Extended Data Fig. 7: Sensitivity analyses for bile acid-associated cell type composition, goblet maturation, and epithelial stress programs. (a)** Correlation of ulcerative colitis (UC) disease activity and host-derived BA cluster a in the scRNA sequencing cohort with Spearman / Pearson coefficients and p-values. (**b**) Selected biopsy-level cell type composition correlates of host-derived BA cluster a. Showing abundances as percent of indicated parent population, plotted against cluster a clr values. Each data point represents one biopsy. Annotations report patient-mean Spearman correlation and biopsy-level random-effects regression coefficient adjusted for Mayo score. (**c**) Biopsy-level random-effects sensitivity analysis of the association between host-derived BA Cluster a and goblet/secretory maturation-state composition. Each data point shows the mayo-adjusted correlation coefficient for cluster a, with patient included as a random intercept. Error bars indicate 95% confidence intervals; asterisks indicate nominal P values ($p<0.05$). (**d**) Reactome pathway GSEA of patient-level pseudobulk expression across the goblet maturation states. Genes were ranked by their association with BA cluster a clr. Showing positive Reactome pathways with FDR < 0.05, ranked by normalized enrichment score (NES); bubble size indicates $-\log_{10}$FDR values. (**e**) Reactome pathway GSEA of CDCA-treated patient-derived colon organoids. Genes were ranked by differential expression in CDCA versus vehicle-treated organoids. Showing positive Reactome pathways with FDR < 0.05, ranked by NES. Bubble size indicates $-\log_{10}$FDR 10 values.
